# Co-Therapy with S1P and Heparan Sulfate Derivatives to Restore Endothelial Glycocalyx and Combat Pro-Atherosclerotic Endothelial Dysfunction

**DOI:** 10.1101/2024.11.06.622347

**Authors:** Ronodeep Mitra, Kaleigh Pentland, Svilen Kolev, Matthew Eden, Erel Levine, Jessica M. Oakes, Eno E. Ebong

**Author notes:** These authors made equal contributions to this work (Ronodeep Mitra, Kaleigh Pentland). Correspondence: Eno E. Ebong, Ph.D., Department of Chemical Engineering, Northeastern University, 805 Columbus Avenue, 221 Interdisciplinary Science & Engineering Complex, Boston, MA 02120.

## Abstract

Endothelial cell (EC) glycocalyx (GCX) shedding due to disturbed blood flow and chemical factors leads to low-density lipoprotein infiltration and reduced nitric oxide synthesis, causing vascular dysfunction and atherosclerosis. This study evaluates a novel therapy combining sphingosine-1-phosphate (S1P) and heparin (heparan sulfate derivative). We hypothesized that heparin/S1P would repair mechanically damaged EC GCX in disturbed flow (DF) regions and restore anti-atherosclerotic mechanotransduction function, addressing cardiovascular disease. We used a parallel-plate flow chamber to simulate flow conditions in vitro and a partial carotid ligation mouse model to mimic DF in vivo. Heparin and albumin-bound S1P were administered to assess their reparative effects on the endothelial GCX. Immunocytochemistry, fluorescent staining, confocal microscopy, cellular alignment studies, and ultrasound were performed to evaluate EC function and endothelial-dependent vascular function. Barrier functionality was assessed via macrophage uptake. Heparin/S1P mechanism-of-action insights were gained through fluid dynamics simulations and staining of GCX synthesis enzyme as well as S1P receptor. Statistical analyses validated results. In vitro data showed that heparin/S1P therapy improves the function of DF-conditioned ECs by restoring EC GCX and promoting EC alignment and elevated vasodilator eNOS (endothelial-type nitric oxide synthase) expression. The in vivo studies confirmed GCX degradation, increased vessel inflammation and hyperpermeability, and vessel wall thickening in the partially ligated left carotid artery. Heparin/S1P treatment restored GCX in the left carotid artery, enhancing GCX thickness and coverage of the blood vessel wall. This work advances a new approach to regenerating the EC GCX and restoring its function in ECs under DF conditions.

## 1. INTRODUCTION

Atherosclerosis is a cardiovascular disease characterized by plaque development and rupture within arterial walls. Atherosclerosis in the context of coronary artery disease, involving plaque formation in heart arteries, account for 75% of heart attacks and remain the leading cause of death globally and in the United States [1]. Notably, 75% of these deaths occur in low-income and underserved regions [2]. Thus, developing novel, accessible, and cost-effective therapeutics is critical.

Endothelium, located between flowing blood and vessel tissue, facilitates vascular homeostasis [3–7], regulating selective permeability, inflammatory processes, and smooth muscle cell proliferation [3,6]. These functions rely on the endothelial glycocalyx (GCX), a negatively charged, hydrated polysaccharide plexus primarily located on the apical surface of endothelial cells (ECs) [8–10]. The GCX is comprised of glycoproteins, glycolipids, and proteoglycans like syndecan-1 (SDC1) and glypican-1 (GPC1), with glycosaminoglycans (GAGs) such as heparan sulfate (HS), hyaluronic acid, and chondroitin sulfate [3,11–13].

An intact GCX is vital for normal endothelial function, regulating vascular permeability and tone via mechanotransduction from undisturbed, uniform, laminar blood flow [4,5,11,14]. Disturbed blood flow patterns from complex vasculature structures can initiate GCX damage and shedding [3,5], resulting in disrupted biochemical signaling [15], increased vascular permeability and consequent inflammation [3,5,8], and reduced vascular tone [11,16,17]. For instance, GCX shedding promotes leukocyte and platelet adhesion [18–20], and GCX shedding upregulates intercellular adhesion molecule-1 (ICAM-1) and vascular cell adhesion molecule-1 (VCAM-1). This increases leukocyte adhesion and macrophage infiltration, triggering inflammation and macrophage infiltration into vessel walls, which ultimately leads to an inflammatory cascade response [21–23]. Furthermore, a direct consequence of EC GCX shedding is impaired vascular tone, where endothelial nitric oxide synthase (eNOS), a precursor to vasodilator nitric oxide (NO), decreases in activity, which reduces NO bioavailability and causes vasoconstriction [10,24,25]. Such impairment is linked to diabetes and hypertension, causing insulin resistance and high blood pressure, respectively [26,27]. Additionally, atherosclerosis onset is often attributed to non-uniform blood flow patterns [3,11,12], causing GCX degradation and subsequent endothelial dysfunction [17]. Thus, targeting the GCX is a crucial first step for preventing and treating cardiovascular disease.

We previously showed that a combined treatment of exogenous HS and sphingosine-1-phosphate (S1P) repaired GCX structure and restored vasculoprotective EC-to-EC communication function in rat fat pad ECs that previously experienced enzymatically induced damage to their GCX and intercellular communication interruption [28]. For reference, HS is the predominant GAG component of the GCX on ECs (>70%) [29]. S1P is a bioactive lipid that stabilizes the vascular system and GCX by enhancing intercellular junction strength via zonula occluden-1 (ZO-1) and connexin, thereby regulating transendothelial permeability [28,30–36]. Since the previous studies conducted in rat fat pad ECs were promising, we sought to further develop and evaluate the restorative potential of this combination therapy to advance its journey toward clinical use.

In this study, we tested the hypothesis that derivatives of HS and S1P can effectively repair damaged GCX structurally and functionally in the initial stages of atherosclerosis in both *in vitro* and *in vivo* conditions. We modified the co-treatment with a more affordable and commercially available derivative to exogenous HS, heparin sodium. Heparin is a widely used anticoagulant that protects and synthesizes endothelial GCX [37]. Specifically, heparin has been indicated to prevent and treat blood clots in patients at risk for thrombotic events and during certain medical procedures [38,39]. It has also been shown to inhibit the shedding of endothelial GCX components into the circulation in disease-free rats [40]. A naturally occurring chaperone, albumin, was utilized to stabilize S1P in an *in vitro* and *in vivo* setting [41,42]. The albumin chaperone bound to S1P improves S1P stability and bioactivity [33].

This new and modified co-treatment was utilized in more relevant *in vitro* and *in vivo* endothelial dysfunction models. One model was achieved by using human coronary artery endothelial cells (HCAECs) cultured *in vitro* using a parallel-plate flow chamber in which disturbed flow (DF) was generated adjacent to undisturbed, uniform, laminar blood flow [11,16]. An *in vivo* model implemented a partial ligation surgery of the left carotid artery (LCA) in a murine model [3,5,43]. Both the *in vitro* and *in vivo* models contained neighboring atherosclerotic-prone DF regions and atheroprotective uniform flow (UF) regions. Upon administration of the co-treatment of albumin-bound S1P (*in vitro* and *in vivo*: 1.5 uM) and heparin (*in vitro*: 2 U/mL; *in vivo*: 500 U/kg) it was determined whether, in early atherosclerosis, GCX damage could be reversed and proper endothelial function (eNOS expression, minimized vascular remodeling, and blocked inflammatory cell infiltration) could be restored. To do this, we assessed endothelial functions including barrier function and vascular tone. We also investigated the co-treatment’s potential mechanism(s) of action, including the possibility of a mechanical mechanism. Finally, we probed the possibility of a biochemical mechanism, either involving targeting of an S1P receptor, or involving chemical synthesis via upregulation of exostosin-like glycosyltransferase-3 (*EXTL3*) that synthesizes HS in the Golgi.

Our study described herein provides further evidence of the potential of HS- and S1P-derived co-treatment to effectively reverse GCX damage and restore proper endothelial function. Moreover, our investigation elucidates crucial underlying mechanisms governing its therapeutic effect. These findings are pivotal for advancing the translation of this treatment into clinical practice, offering a promising avenue for improving cardiovascular disease management and ultimately enhancing patient outcomes.

### 2. MATERIALS & METHODS

#### 2.1 Overview

Studies conducted in this paper involve both *in vitro* and *in vivo* experiments that utilize unique complex models, which incorporate two flow regions. There is an atheroprone DF region characterized by recirculation and stagnation. In addition, there is an atheroprotective UF region characterized by unidirectional laminar shear stress. The *in vitro* and *in vivo* effects of DF and UF on endothelial GCX expression and endothelial mechanotransduction were assessed, with the expectation of impaired GCX and endothelial function in DF compared to UF conditions. Co-treatment of albumin-bound S1P and heparin were administered *in vitro* and *in vivo* to test therapeutic efficacy in reversing GCX damage and restoring proper endothelial function, including barrier function and vascular tone. The mechanism of action of the co-treatment was assessed, focusing on mechanisms involving mechanical factors (fluid and particle simulation) and chemical factors (receptor targeting and biosynthesis of glycosaminoglycans).

#### 2.2 Availability of Data

The authors declare that all supporting data are available within the article and its online supplementary files.

#### 2.3 Disturbed Flow (DF) Models

Both *in vitro* and *in vivo* atherosclerotic-prone DF models were used in this study to accelerate endothelial dysfunction, a hallmark and initiator of atherosclerotic disease [17]. Specifically, a parallel-plate flow chamber cell culture model [11,16] and a partial ligation of left carotid artery (LCA) murine model [3,5,44] were used to induce acute DF patterns *in vitro* and *in vivo*, respectively. Both models also have adjacent atheroprotective UF patterns to mitigate sample prep and the number of cell culture and animal samples.

##### Parallel-Plate Flow Chamber (Shear Stress Apparatus) In Vitro

Human coronary arterial ECs (HCAECs) purchased from PromoCell (Heidelberg, Germany) were cultured between passages 4 and 8 in PromoCell Endothelial Cell Growth Medium MV2 supplemented with growth factors, fetal calf serum, and antibiotic penicillin streptomycin. ECs were grown and maintained in a sterile humidified incubator at 37 °C with 5% CO2. For shear stress experimental use, HCAECs were seeded at a density between 21,000 – 31,000 cells/cm^2^ on human protein-derived fibronectin (Gibco^TM^) (60 µg/mL) coated glass coverslips. HCAECs were then transferred to the flow chamber system with PromoCell Endothelial Cell Growth Medium MV2 supplemented with 0.5% bovine serum albumin (BSA) circulating through the chamber. This exposed the HCAECs to dynamic shear stress [11,16,44]. The chamber was designed to simulate atheroprone DF patterns upstream, characterized by fluid recirculation, stagnation, and directional gradient, with an eventual downstream recovery to unidirectional 12 dynes/cm^2^ atheroprotective UF patterns. 12-hour flow condition HCAECs were compared to HCAECs at 0-hour baseline conditions (these cells were acclimated in BSA-containing experimental medium for 30 minutes) and HAECs left in static conditions for concurrent 12 hours. Therefore, three different cell culture groups were examined: 1) 0-hour static (baseline EC behavior to which data was normalized); 2) 12-hour static; and 3) 12-hour flow (includes both UF and DF conditions).

##### Partial Carotid Ligation Murine Model In Vivo

All animal studies were conducted under protocol 23– 0716R, approved by the Northeastern University Institutional Animal Care and Use Committee (NU-IACUC). 10–12-week-old male C57Bl/6 mice were obtained from Jackson Laboratories and were housed in Northeastern University’s animal facility in a standard laboratory environment with an ambient temperature of 22-25℃, humidity of 55-65%, and a 12/12 hour light/dark cycle. Animals were acclimated to these conditions and fed a regular chow diet and water *ad libitum* for at least 1 week as recommended by NU-IACUC. For experiments, all mice were subjected to partial ligation surgery of their left carotid artery (LCA) to induce acute disturbed blood flow patterns and accelerate endothelial dysfunction and GCX degradation, as previously described [3,5,43–45]. The right carotid artery (RCA) was left intact to provide a reference vessel in each mouse. A single subcutaneous injection of Meloxicam (5 mg/mL) was administered immediately before surgery for pain relief. Upon post-surgery recovery, mice were returned to the animal care facility, where standard chow diet and water were provided *ad libitum*. Mice were monitored daily to ensure proper recovery. For experiments, the groups examined were: 1) ligated LCA (for DF conditions); versus 2) non-ligated RCA (for UF conditions).

#### 2.4 HS and Albumin-Bound S1P Co-Treatment Administration

##### In Vitro HS and Albumin-Bound S1P Co-Treatment Administration

HCAECs were either not treated or exposed to co-treatment for the 12-hour duration of the flow, which included heparin (2 U/mL; Sigma Aldrich) and albumin-bound S1P (1.5 uM; Avanti Polar Lipids) supplemented into MV2 media with 0.5% BSA. Albumin-bound S1P was created using the protocol suggested by Avanti Polar Lipids. Briefly, C17-S1P (1 mg; Avanti Polar Lipids) was resuspended in 13.4 mL of methanol (organic solvent) for 12 hours and dried under vacuum. Fresh albumin-bound S1P solution was made before each experiment by adding 0.4% solution of PBS-Albumin (fatty acid free; Sigma) to dried S1P and subjected to a quick bath sonication.

##### In Vivo HS and Albumin-Bound S1P Co-Treatment Administration

5 days post-LCA ligation surgery (as described above), the mice were either not treated or administered vehicle only (saline) or co-treatment (albumin-bound S1P combined with heparin). A 15 μmol/L stock solution of C17-S1P (Avanti Polar Lipids) was prepared with 4% mouse serum albumin in sterile saline, as previously described [33]. A mixture of 1.5 μmol/L albumin-bound S1P (∼ 150 uL) and 500 U/kg Heparin (∼ 10 – 12 uL) was created and delivered to the mouse via intravenous retro-orbital injection with a sterile 31-gauge ultra-fine needle U100 insulin syringe. Due to the approximate 15-minute half-life of albumin-bound S1P and the approximate 30-minute half-life of heparin [33,46], at 30 minutes after therapy injection, the mice were imaged via ultrasound or sacrificed to collect their vessel tissue for further processing.

#### 2.5 Staining of Endothelial GCX GAGs, Core Proteins, and S1P Receptor – 1

Immediately after exposure to different stimulus conditions and treatments, HCAECs and mouse tissue were stained to probe different biomarkers and assess response to heparin and albumin-bound S1P co-treatment vs. no treatment conditions. These biomarkers included whole GCX, HS (most abundant GAG of the GCX), GCX core protein syndecan-1 (SDC1; binds to HS and chondroitin sulfate), and GCX core protein glypican-1 (GPC1; binds to HS). We also stained the S1P receptor 1 (S1PR1; specifically present on endothelial cells).

##### In Vitro Methods

Descriptions of methods applied to HCAECs are described in **Table I**.

**Table I.** Staining Protocols for GCX Components and S1PR1 (in vitro). Newly mentioned abbreviations are as follows: Ab – antibody, AF – Alexa Fluor, BS – blocking serum, BSA – bovine serum albumin, GA - glutaraldehyde, PBS – phosphate buffered saline, PFA – paraformaldehyde, WGA – wheat germ agglutinin. Please note that the protocols outlined in this table only apply in vitro and do not apply in vivo.

| Biomarker | Fixative | Permeabilization | Blocking Solution (BS) | Primary Antibody (Ab) | Secondary Ab | Tertiary Ab | Mounting Medium |
| --- | --- | --- | --- | --- | --- | --- | --- |
| Heparan Sulfate | 2% PFA + 0.1% GA (30 minutes) | n/a | 2% goat serum in PBS (1 hour) | 1:100 HS antibody, 10e4 epitope in BS, 4 °C (2-3 days) | 1:400 AF488 goat anti-mouse IgM in BS (30 minutes) | n/a | VectaShield (Vector Labs) anti-fade mounting medium with 4',6-diamidino-2-phenylindole (DAPI) |
| Whole GCX (WGA) |  |  | 3% BSA in PBS (1 hour) | 1:200 Biotinylated WGA Lectin in BS (1 hour) | 1:500 AF488 or AF647 streptavidin in BS (30 minutes) |  |  |
| Syndecan-1 |  | 0.1% Triton in PBS (10 minutes) | 5% goat serum in PBS (1 hour) | 1:100 SDC1 (CD138) anti-rabbit monoclonal in BS, 4 °C (2-3 days) | 1:400 AF488 goat anti-rabbit IgG (1:400) in BS (2 hours) |  |  |
| Glypican-1 | 2% PFA (30 minutes) | 0.1% Triton in PBS (5 minutes) | 5% BSA in PBS (1 hour) | 1:100 GPC1 mouse monoclonal antibody in BS, 4 °C (3 days) | 1:400 Biotinylated goat anti-mouse in BS (1 hour) | 1:400 AF488 Streptavidin in PBS (1 hour) |  |
| S1PR1 | 2% PFA (30 minutes) | 0.1% Triton in PBS (5 minutes) | 5% BSA in PBS (1 hour) | 1:100 S1PR1 mouse monoclonal antibody in BS (3 days) | 1:400 Biotinylated goat anti-mouse in BS (1 hour) | 1:400 AF488 Streptavidin in PBS (1.5 hours) |  |

##### In Vivo Methods

Prior to processing the mouse tissue, an extensive process of LCA and RCA tissue dissection, preservation, and cryosectioning was implemented, as previously described [3,5].

Subsequently, to assess GCX integrity *in situ*, immunohistochemical analysis was performed on sectioned LCA and RCA using WGA lectin and SDC1 antibody, according to a prior published protocol [3].

Similar approaches, with minor modifications, were used to stain GPC1. Briefly, previously perfusion fixed tissue samples on glass slides were post-fixed in 4% PFA for 10 minutes and then washed thoroughly with PBS. Tissue samples were then permeabilized for 10 min with 0.3% Triton X-100 diluted in PBS. Subsequently, antigen retrieval was achieved by heating samples in 10 mM sodium citrate at a pH of 6.0 in PBS (for 10 min) and allowing them to equilibrate to room temperature. Next, endogenous peroxidase blocking was achieved by using a 1% hydrogen peroxide solution diluted in deionized water for 30 min with rocking. Tissue samples were then introduced to further blocking solution (10% goat serum and 0.3% Triton) for 1 hour in a humidified condition. Slides were then incubated with polyclonal GPC1 antibody (Invitrogen; 1:200 in blocking solution) for 3 days. Following primary antibody incubation, tissue samples were exposed to a goat anti-rabbit secondary antibody conjugated to horse radish peroxidase (1:250 dilution in PBS) and incubated for 1 hour in a humidified condition at room temperature. Tissue samples were subsequently washed in a 0.1% Tween 20 (Sigma-Aldrich Co.) solution in PBS. A TSA Cyanine 3 amplification system kit from Akoya Biosciences (Part No. NEL704A001KT) was then used to incubate tissue samples for 5 min. Finally, the slides were washed again in 0.1% Tween 20 in PBS.

Stained tissue samples were mounted and sealed using VectaShield anti-fade mounting medium with DAPI for cell nuclei staining (VectorLabs, part number H1200).

#### 2.6 Assessing HS and Albumin-bound S1P Co-Treatment Effect on Endothelial Cell and Vascular Function

##### General Methods

Vascular tone regulation and remodeling as well as barrier function were monitored to assess the co-treatment’s ability to reverse endothelial dysfunction *in vitro* and *in vivo*. To assess vascular tone regulation *in vitro*, HCAECs were stained for serine 1177 phosphorylated endothelial nitric oxide synthase (p-eNOS; active form) and total eNOS. Furthermore, remodeling of cultured HCAEC morphology was assessed (round versus elongated in the direction of flow). To assess vascular tone and remodeling *in vivo*, periodic ultrasound data was collected to determine diastolic and systolic lumen diameter along with vessel wall thickness of the LCA and RCA. Macrophage uptake via CD68 was assessed for hyperpermeability *in vivo*. Further details of studies are described below:

##### Monitoring In Vitro Cellular Remodeling, As Indicated by EC Morphology

Cell orientation and alignment analysis was done using a custom Python script designed to analyze brightfield images of cells in the area of interest within a flow-conditioned HCAEC monolayer. 4 regions were imaged for each condition, and each region was analyzed to obtain an alignment score as follows. First, images were run through an automatic local thresholding using the Otsu method with a local radius of 100, and morphological closing (erosion + dilation) was applied. Isolated connected objects were labeled as unique objects, and all objects were filtered for reasonable size determined by area to exclude small specs and background regions. This yielded a mask that identified the cells as blobs. An ellipse was fitted to each cell blob to return the cells’ center, length, width, and angle of orientation. For each sample image, 500-1000 cells were identified and measured. Manual inspection was performed to ensure quality segmentation and ellipse fit. Subsequently, to measure the alignment of cells in each experiment, identified cells were filtered to exclude cells that were not elongated beyond an aspect ratio of 2. The angle of orientation was adjusted to accommodate cell orientation, which is bidirectional. Alignment was measured in two ways: 1) Object Orientation Parameter (OOP) and 2) median pairwise alignment (MPA). The OOP score is computed as OOP = 2cos^2^(Ɵ) – 1, where Ɵ is the orientation angle. OOP ranges between -1 and +1, where an OOP = -1 indicates alignment perpendicular to the flow direction, while an OOP = +1 indicates alignment parallel to the direction of flow [47]. MPA measures alignment of cells to each other and is computed as the median of the absolute value of pairwise dot product. MPA for a large number of pairs ranges from 0.5 to 1. A score of 1 indicates all cells are perfectly aligned and oriented in the same direction while a score of 0.5 indicates random orientations.

##### Monitoring EC Control of Vascular Remodeling, As Indicated by In Vitro Expression of Vascular Tone Regulators Including Active (phosphorylated) eNOS (p-eNOS) and Total eNOS

HCAECs were first fixed in a 4% PFA solution for 20 minutes. Samples were then permeabilized in 0.5% triton in PBS solution for 15 minutes and subsequently blocked in 10% goat serum in PBS for 1 hour at room temperature. The coverslips were incubated with their respective primary antibody at a 1:500 dilution in blocking solution in a humidified chamber for 3 days at 4℃. The antibody used for the p-eNOS targeted eNOS that was phosphorylated at serine 1177, which represents the active form of eNOS. The antibody that was used to target total eNOS was the D9A5L version. After primary antibody incubation and thorough washing, samples were incubated with Alexa Fluor 488 labeled goat anti-rabbit secondary antibody (1:200) in PBS for 1 hour in dark, room temperature conditions. Once staining was completed, samples were mounted and sealed using VectaShield anti-fade mounting medium with DAPI for cell nuclei staining (VectorLabs, H1200).

##### Monitoring Vascular Remodeling Via High-Resolution Ultrasound Measurements

Blood vessel diameter and wall thickness were measured using ultrasound, which was conducted with Fujifilm Visual Sonics Vevo® 3100 LT with MX700 (29-71 MHz; 10 mm scan depth) and MX550S (25-55 MHz; 15 mm scan depth) transducers. All RCAs and LCAs of mice were periodically imaged on day 0 (before LCA ligation), day 3 (post-ligation), and day 5 (post-ligation and 30 minutes after co-treatment injection). Mice were anesthetized with inhaled isoflurane (1-2.5%), and body temperature was maintained on a heated 38℃ platform. Levels of anesthesia, heart rate, temperature, and respiration were continuously monitored during the imaging session. Pulse-wave Doppler Mode was used to assess blood flow velocity through the carotid artery to ensure partial ligation of the LCA was successful. Pulse-wave Doppler Mode can detect any blood flow disturbances in the vessel or region of bifurcation. *M*-mode was used for vessel dimensions. Parameters that were calculated include vessel wall thickness and diastolic and systolic lumen diameter. All measurements were gated to an electrocardiogram and respirator.

##### Monitoring In Vivo Barrier Functionality, As Indicated by Vessel Wall Infiltration by Macrophages

LCA and RCA infiltration of macrophages were assessed via cluster of differentiation 68 (CD68) staining. The full protocol was previously published [3,5].

#### 2.7 Assessment of HS and Albumin-Bound S1P Co-Treatment Mechanisms of Action

##### General Methods

Understanding the mechanism of action of the co-treatment was achieved in two ways:

1. by determining the residence time of the therapy in DF, via fluid and particle simulations, 2) by determining if the co-treatment acts at all via the S1P receptor pathway, and 3) by determining whether there is active nascent proteoglycan synthesis via exostosin-like glycosyltransferase-3 (*EXTL3*). Detailed protocols are described below:

##### Simulations to Assess a Potential Mechanical Mechanism of Action

The fluid flow in the millifluidic device was modeled as an incompressible Newtonian fluid with a density of 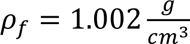 and dynamic viscosity of μ = 0.862 *mPa* · *s*. The steady-state velocity and pressure fields in the millifluidic device (Ω) were governed by the Navier-Stokes equations: 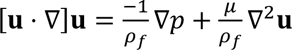, where **u** represents the fluid velocity vector, *p* represents the pressure, and ρ_*f*_ represents fluid density. The steady-state Navier-Stokes equations were solved using the *simpleFoam* solver through the open-source computational fluid dynamics software OpenFOAM. The millifluidic parallel-plate flow device can hold up to four glass cover-slips to increase experimental throughput. The flow chamber consists of 4 individual “steps” to ensure each cover-slip with HCAECs is exposed to atheroprone DF conditions upstream of atheroprotective UF conditions. Initial simulations demonstrated identical flow fields over each “step” within the millifluidic device. Thus, it was optimal to simulate fluid and particle transport over only one “step”. The computational domain was discretized into 9.9 million orthogonal elements using OpenFOAM’s *blockMesh* utility. The inlet was assigned a constant flux Dirichlet boundary condition of 102 mL/min, the outlet was assigned a zero-pressure Neumann boundary condition, and the walls were assigned a no-slip boundary condition. Following fluid simulations, the transport of co-treatment of albumin bound S1P and heparin was simulated through OpenFOAM’s Lagrangian particle tracking solver, *particleFoam*. Particles were assumed not to influence the flow field nor interact with other particles. Particles were also assumed to be well-mixed throughout the medium and thus were initialized uniformly throughout the domain of interest. The particle’s position is updated through the equation 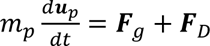 where ***u***_*p*_ is the particle velocity, *m*_*p*_ is the particle mass, ***F***_*g*_ is the gravitation force, and ***F***_*D*_ is the drag force. Based on experimental data obtained via dynamic light scattering (*see Supplemental Figure 1)*, all particles were assigned to have a diameter of *d*_*p*_ = 730 ρ*m* with a density of 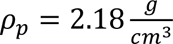. Furthermore, all particles were assumed to be spherical. When a particle’s center was within one radius of the wall of the microfluidic device, it was assumed to stick to the wall and was no longer tracked.

##### Assays Performed to Assess Potential Mechanism of Action Involving the S1P Receptor Pathway

See in vitro methods described in Section 2.5 and Table 1.

##### Experiments Performed to Assess Potential Mechanism of Action Involving Biochemical Synthesis of GCX Replacement Components

Exostosin-like glycosyltransferase-3 (*EXTL3*) was stained since it plays an important role in the chain polymerization of HS and heparin via the Golgi apparatus [48]. Anticipating that the heparin and albumin-bound S1P co-treatment could exert their therapeutic activity through upregulation of *EXTL3* activity within vascular ECs, for mice experiments we stained LCA and RCA tissue for *EXTL3*. Fixed tissue samples were treated with 10% goat blocking serum and 0.3% Triton. Tissue samples were then either incubated with *EXTL3* polyclonal antibody (diluted 1:100 in blocking solution) for 3 days at 4 °C. Following primary antibody incubation, tissue samples were coated with 1:250 dilution in PBS of HRP-conjugated goat anti-rabbit antibody for 1 hour. TSA Cyanine 3 amplification system kit was then used to incubate the tissue samples for 5 minutes. After washing thoroughly in 0.1% Tween 20 solution, all tissue samples were mounted and sealed using VectaShield anti-fade mounting medium with DAPI for cell nuclei staining (VectorLabs, H1200).

#### 2.8 Confocal Microscopy Imaging

All samples (except where otherwise indicated above) were imaged using a LSM 800 confocal microscope (Carl Zeiss Meditec AG, Jena, Germany) housed in the Institute for Chemical Imaging of Living Systems (CILS) core facility at Northeastern University. HCAEC samples were imaged at 63x (with an oil immersion lens) magnification with z-stack, while murine LCA and RCA samples were imaged at 20x tile scans and 40x magnification. An excitation wavelength of 360 nm was used to capture DAPI-labeled cell nuclei, and a 554 nm wavelength showed the Cy3 fluorescently amplified labels of biomarkers in mice tissue samples. Additionally, 488 nm wavelength was used to visualize elastin auto-fluorescence (vessel wall) in mice carotid artery samples and to capture fluorescently tagged biomarkers of interest in HCAECs.

#### 2.9 Image Analysis and Data Acquisition

For *in vitro* experiments, to determine overall expression levels of HS in HCAECs, ImageJ was used to calculate the mean fluorescence intensity (MFI) from 63x *en face* images [11]. Percent area fractions were computed to quantify WGA, p-eNOS, total eNOS, GPC1, SDC1, and S1PR1 biomarkers of interest. Three different 63x z-stack images were taken from each sample and were averaged. All data from each run was normalized to their respective zero-hour experimental group. Phase contrast images were also captured and analyzed using a customized Python script to quantify cell alignment from six different experimental runs from each HCAEC treatment group.

For *in vivo* experiments, GCX coverage and thickness probed via labeling WGA, SDC1, and GPC1 in the LCA and RCA, as well as barrier function probed by labeling macrophage uptake within vessel walls, were imaged and analyzed via ImageJ as previously described [3,5]. *EXTL3* images were analyzed by determining the percent area fraction of expression within the endothelium. For all biomarkers *in vivo*, three serial tissue rings were quantified and then averaged for one data point per LCA or RCA of each animal. Quantifications of various biomarkers were normalized to their RCA counterpart. Lastly, ultrasound image data was collected and analyzed via software Vevo Lab 5.6.1.

#### 2.10 Statistics

All data sets (raw and normalized data) were represented as mean ± standard error of the mean (SEM) with a minimum of 4 independent experiments per condition, upon which significance testing in the accompanying figures were conducted. Comparisons among more than two groups were either conducted using Ordinary one-way ANOVA with Tukey’s multiple comparisons test, or two-way ANOVA with Tukey’s multiple comparisons test, using GraphPad Prism software (version 9.5.0). The results were also plotted using GraphPad Prism, where an α-value of 0.05 and a 95% confidence interval were used for statistical significance. In data figures, asterisks indicate statistical significance of the data, as follows: *p < 0.05, **p < 0.01, ***p < 0.001, and ****p <0.0001.

### 3. RESULTS

#### 3.1 Co-Treatment Impact on EC GCX Integrity

This portion of the results section highlights the impact that co-treatment of albumin-bound S1P and heparin has on EC GCX integrity *in vitro* and *in vivo*. As a reminder, the *in vitro* markers of the EC GCX include WGA, HS, SDC1, and GPC1. The *in vivo* markers of the EC GCX include WGA, SDC1, and GPC, with HS being omitted because the HS antibody is made in mouse and produces significant non-specific staining. The results obtained are described below.

##### Co-Treatment of Albumin-bound S1P and Heparin increases WGA expression in DF-atheroprone region in vitro and in vivo

Figure 1 displays representative images of HCAECs stained for WGA (labeled as green) across experimental groups and data corroborating previous studies performed using parallel-plate flow chambers [11,16,44]. Specifically, we assessed 12-hour flow-induced HCAEC coverage by GCX labeled using WGA lectin. We normalized the data to a 0-hour baseline within each experiment to account for variations. Under static conditions, untreated HCAECs showed a normalized WGA expression of 1.03 ± 0.05. In DF conditions, this decreased to 0.70 ± 0.09, and in UF conditions, it increased slightly to 1.13 ± 0.04. DF conditions showed a significant 32% decrease compared to static, and a 38% decrease compared to UF conditions. No significant difference was observed between static and UF conditions. Upon exposure to co-treatment, WGA-labeled GCX expression significantly increased. In DF conditions, co-treatment resulted in a normalized WGA expression of 3.96 ± 0.63, compared to 0.70 ± 0.09 for untreated DF conditions. Similarly, in UF conditions, co-treatment showed a normalized expression of 3.39 ± 0.66, compared to 1.13 ± 0.04 for untreated UF conditions. In static conditions, co-treated HCAECs expressed 3.39 ± 0.66, compared to 1.03 ± 0.05 for untreated conditions.

**Figure 1.**
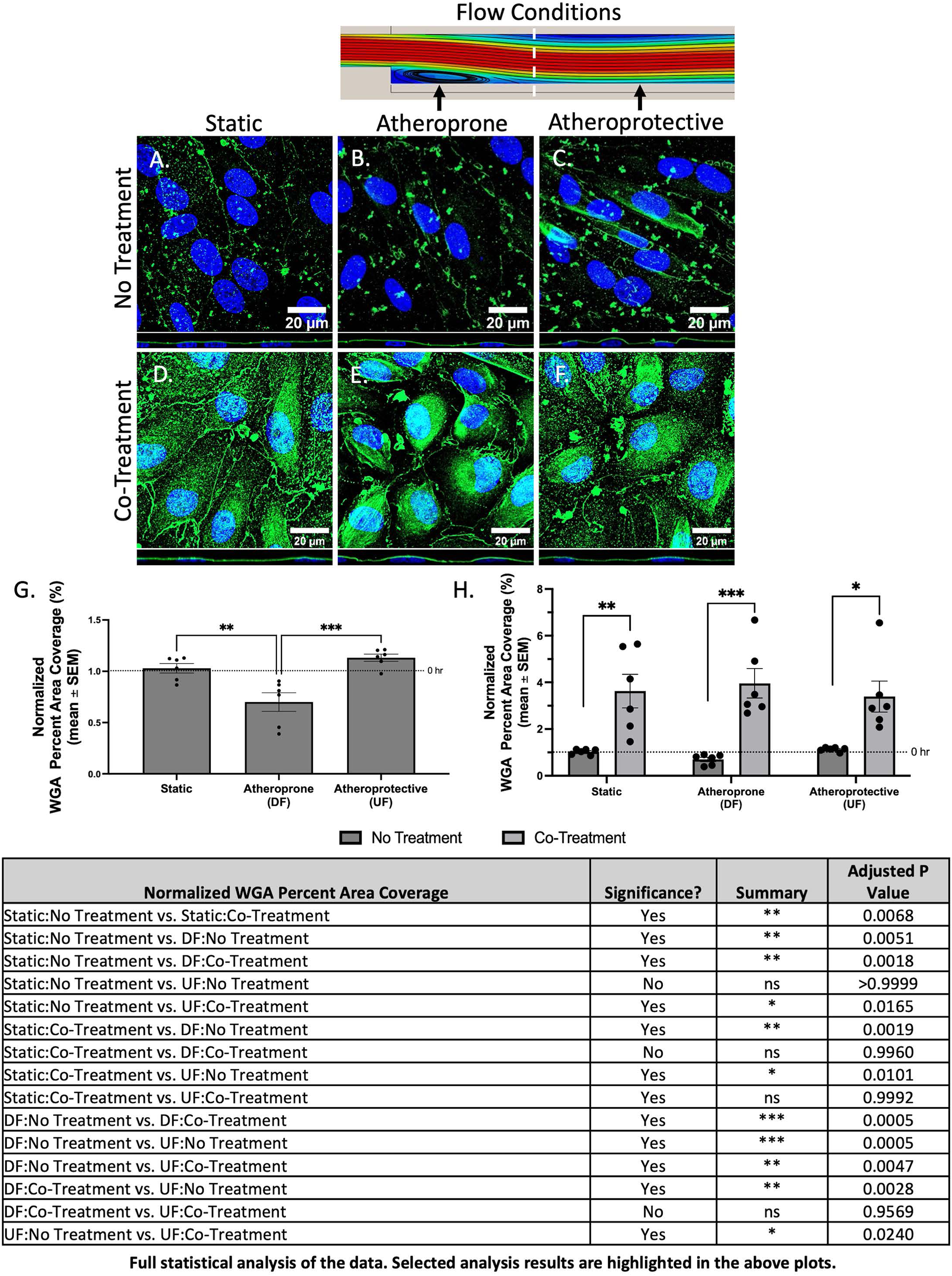
Effect of co-treatment on WGA expression in HCAECs after 12 hours of flow at 12 dynes/cm^2^. Shown are 63x representative images of HCAECs in each experimental group. Fluorescent staining of wheat germ agglutinin (WGA; green), a lectin that stains multiple components of the GCX, was used to provide a comprehensive view to assess the percent area coverage of the whole structural GCX. WGA expression increases in all conditions when co-treatment is administered with HCAECs. Blue represents cell nuclei labeled with DAPI. The scale bar is 20 μm. HCAECs in the no treatment group under A) static conditions, B) in atheroprone conditions, and C) under atheroprotective conditions. HCAECs in the co-treatment group under D) static conditions, E) atheroprone conditions, and F) atheroprotective conditions. Graphs show the mean ± SEM of percent area coverage of WGA staining normalized to the control (0 hr static condition) for G) no treatment group only and H) all cohorts (n = 6). Statistical analysis was performed using a one or two-way ANOVA. Significance is denoted by asterisks: *p<.05, **p<.01, ***p<.001, and ****p<.0001. *Note: Supplemental Figure 2 contains results that show WGA expression in HCAECs that were treated with albumin-bound S1P or heparin separately*.

Figure 2 presents *in vivo* data validating the *in vitro* (Figure 1) findings. We previously published that in untreated conditions the WGA-labeled GCX on the non-ligated RCA in C57BL/6 male mice expresses a WGA fluorescence pattern that spans 76.3 ± 10.2% of the inner blood vessel wall [44]. In the present study, the WGA (labeled as red) fluorescence pattern was found to present even higher RCA coverage of 86.6 ± 1.40% with vehicle treatment, 89.2 ± 1.03% with co-treatment of albumin-bound S1P and Heparin, and 92.2 ± 0.64% in untreated conditions. The ligated LCA, normalized to the RCA per condition, showed a normalized vessel coverage of 0.65 ± 0.03 in untreated conditions, 0.65 ± 0.05 with vehicle-only, and 0.95 ± 0.02 with co-treatment, indicating that albumin-S1P/heparin results in a significant 46% increase compared to untreated and 48% increase compared to vehicle-only conditions. Regarding GCX thickness, on the RCA wall, untreated conditions measured 1.45 ± 0.083 μm, vehicle-only 1.65 ± 0.079 μm, and co-treatment 1.56 ± 0.063 μm, with no significant differences between conditions. In the ligated LCA, thickness ranged from 0.90 μm to 2.05 μm across all conditions. When the data was normalized to the RCA, the readout is 0.74 ± 0.02 for untreated, 0.75 ± 0.02 for vehicle-only, and 1.08 ± 0.04 for co-treatment conditions. Co-treatment induced a significant 46% increase compared to untreated and 44% increase compared to vehicle-only conditions, returning GCX thickness to healthy RCA levels.

**Figure 2.**
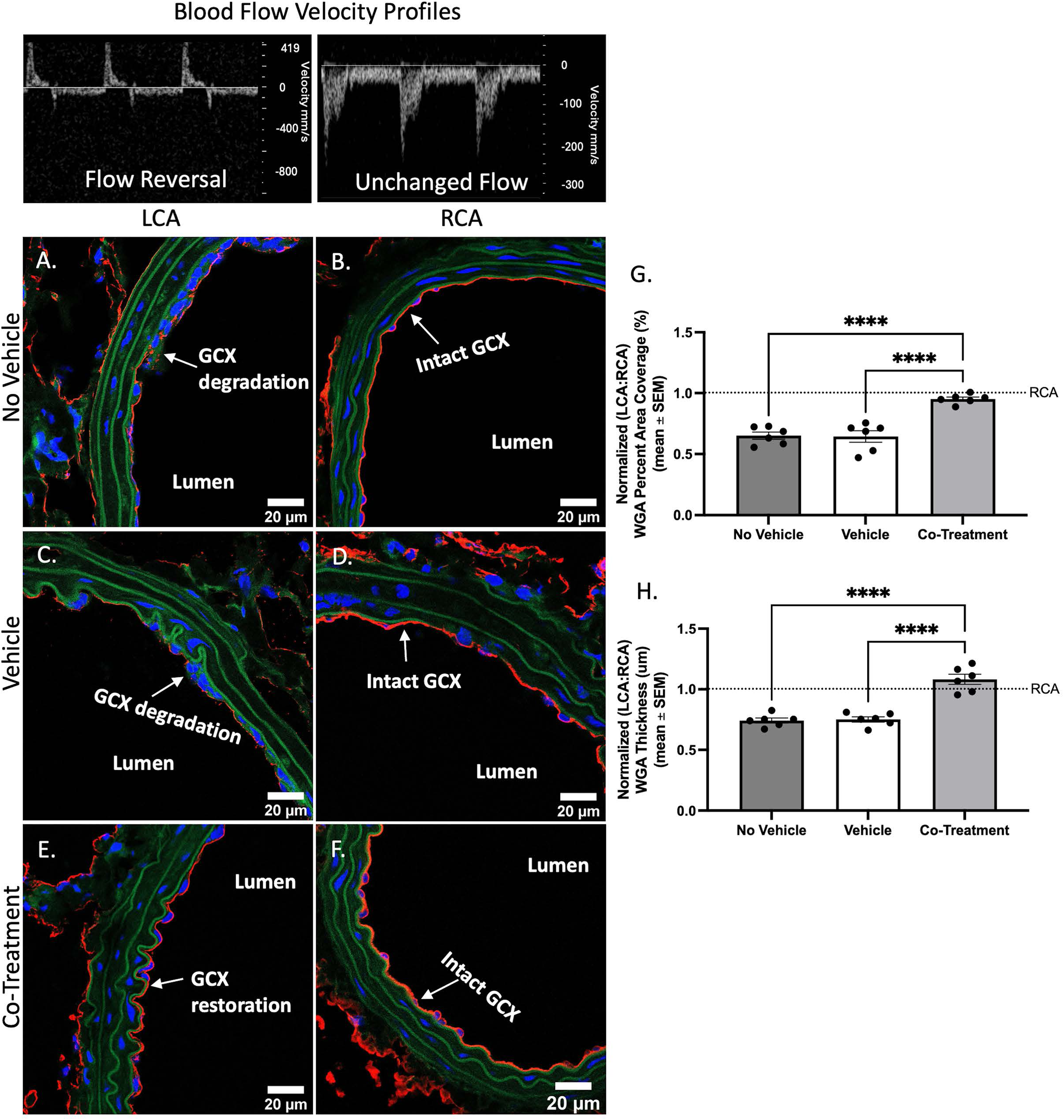
Effect of co-treatment on WGA expression in murine carotid arteries. Shown are 40x representative images of carotid arteries from each cohort of mice. Fluorescent staining of wheat germ agglutinin (WGA; red), a lectin that stains multiple components of the GCX, was used to provide a comprehensive view of the whole structural GCX on the luminal side of the carotid arteries. The percent area coverage and the thickness of WGA increased following a single dose of the co-treatment administered via retro-orbital injection. Blue represents cell nuclei labeled with DAPI. Green represents elastin. The scale bar is 20 μm. A) Ligated LCA from the no vehicle group. B) Control RCA from the no vehicle group. C) Ligated LCA from the vehicle only group. D) Control RCA in the vehicle only group. E) Ligated LCA from the co-treatment group. F) Control RCA from the co-treatment group. G) Graph shows the mean ± SEM of percent area coverage of WGA staining normalized to the control (RCA of each mouse; n = 6). H) Graph shows the mean ± SEM of WGA transversal thickness normalized to the control (RCA of each mouse; n = 6). Statistical analysis was performed using a one-way ANOVA. Significance is denoted by asterisks: *p<.05, **p<.01, ***p<.001, and ****p<.0001.

##### Co-Treatment of Albumin-bound S1P and Heparin increases HS expression in DF-atheroprone region in vitro

Figure 3 shows representative images of stained HS component of the GCX on culture HCAECs, and analysis using mean fluorescence intensity (MFI) quantification as we have done in our prior published work [11]. MFI data was normalized to a 0-hour baseline within each experiment to account for variations. Normalized HS MFI in 12-hour static HCAECs was 1.17 ± 0.08, 12-hour DF HCAECs was 1.11 ± 0.08, and 12-hour UF HCAECs was statistically significantly high 1.57 ± 0.06. After establishing HCAEC HS MFI expression in untreated static, DF, and UF conditions, HCAECs were exposed to co-treatment of albumin-bound S1P and heparin to determine the efficacy of HS GAG restoration. The co-treated HCAEC HS MFI showed a normalized level of 1.11 ± 0.08 in static conditions, 5.11 ± 0.814 in DF conditions, and 3.92 ± 0.534 in UF conditions. It is notable that the co-treatment effects were statistically significant in DF and UF conditions but not statistically significant in 12-hour static conditions.

**Figure 3.**
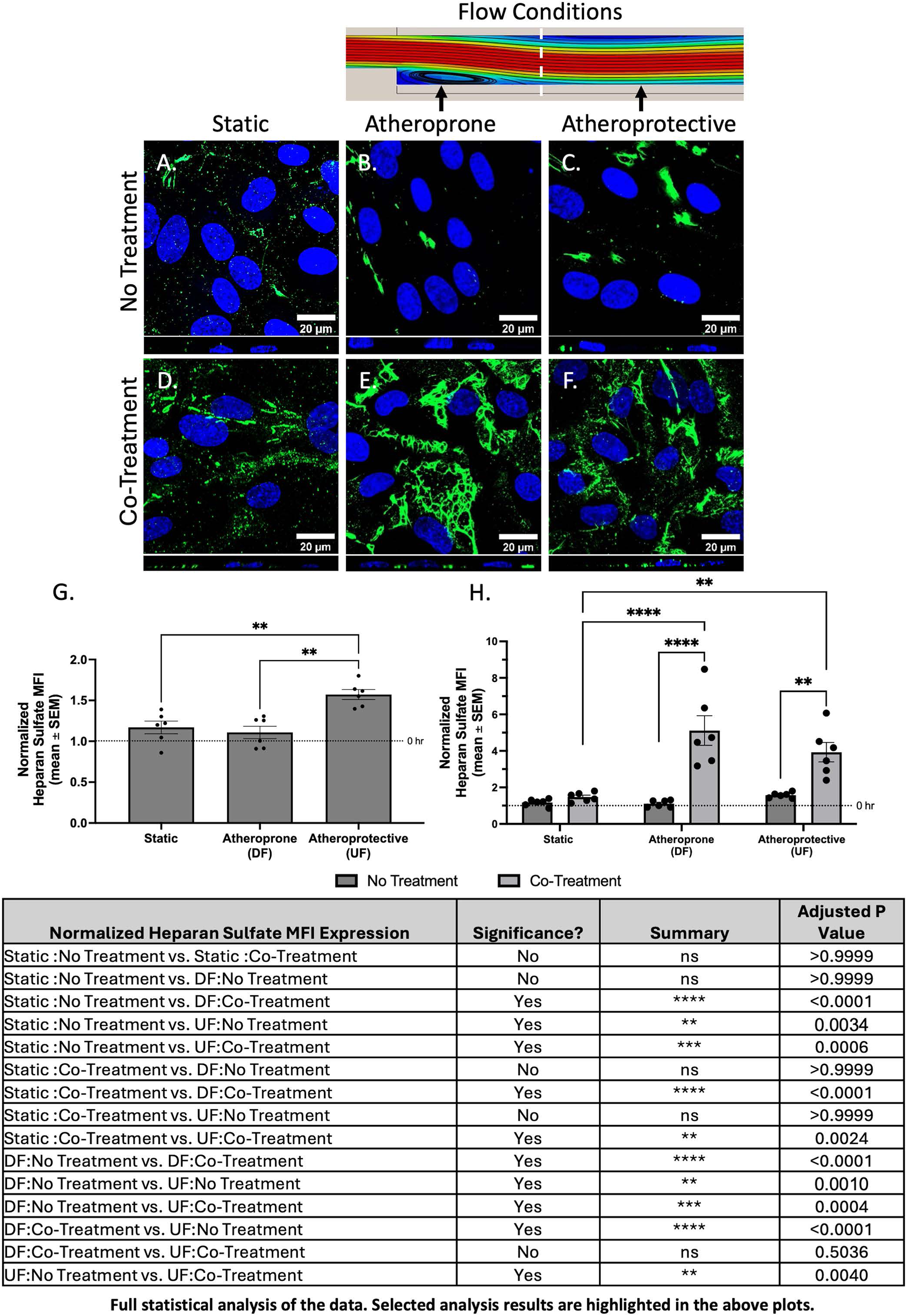
Effect of co-treatment on HS expression in HCAECs after 12 hours of flow at 12 dynes/cm^2^. Shown are 63x representative images of HCAECs in each experimental group. Fluorescent staining of HS (green), a key GAG component of the GCX, increased in both the atheroprone and atheroprotective flow conditions with the co-treatment. Blue represents cell nuclei labeled with DAPI. The scale bar is 20 μm. HCAECs in the no treatment group under A) static conditions, B) atheroprone conditions, and C) atheroprotective conditions. HCAECs in the co-treatment group under D) static conditions, E) atheroprone conditions, and F) atheroprotective conditions. Graphs show the mean ± SEM of the MFI of HS staining normalized to the control (0 hr static condition) for G) no treatment group only and H) for all cohorts (n = 6). Statistical analysis was performed using a one or two-way ANOVA. Significance is denoted by asterisks: *p<.05, **p<.01, ***p<.001, and ****p<.0001. *Note: Supplemental Figure 3 contains results that show HS expression in HCAECs that were treated with albumin-bound S1P or heparin separately*.

##### Co-Treatment of Albumin-bound S1P and Heparin increases core protein SDC1 and GPC1 expression in DF-atheroprone region in vitro and in vivo

To further assess the impact of the co-treatment on the HS component of the EC GCX, we probed the *in vitro* and *in vivo* expression of the transmembrane and cytoskeleton-linked SDC1 and membrane and lipid-raft-linked GPC1 [49]. SDC1 and GPC1 are core GCX proteins that specifically covalently bind HS [49]. Figure 4 shows 63x z-stack representative images of SDC1 on HCAECs *in vitro* in different stimulus environments. Quantifying a full set of image data resulted in the following findings when the data were normalized to 0-hour baseline conditions. In no treatment conditions, normalized SDC1 percent area coverage was observed to be 1.04 ± 0.04 in 12-hour static conditions, 0.75 ± 0.09 in 12-hour DF conditions, and 1.14 ± 0.07 in 12-hour UF conditions, showing that SDC1 was statistically significantly impaired by DF conditions compared to static or UF conditions. In conditions of co-treatment with albumin-S1P and heparin, normalized SDC1 percentage area increased to 1.66 ± 0.20 in 12-hour static conditions, 1.65 ± 0.22 in 12-hour DF conditions, and 1.90 ± 0.275 in 12-hour UF conditions.

**Figure 4.**
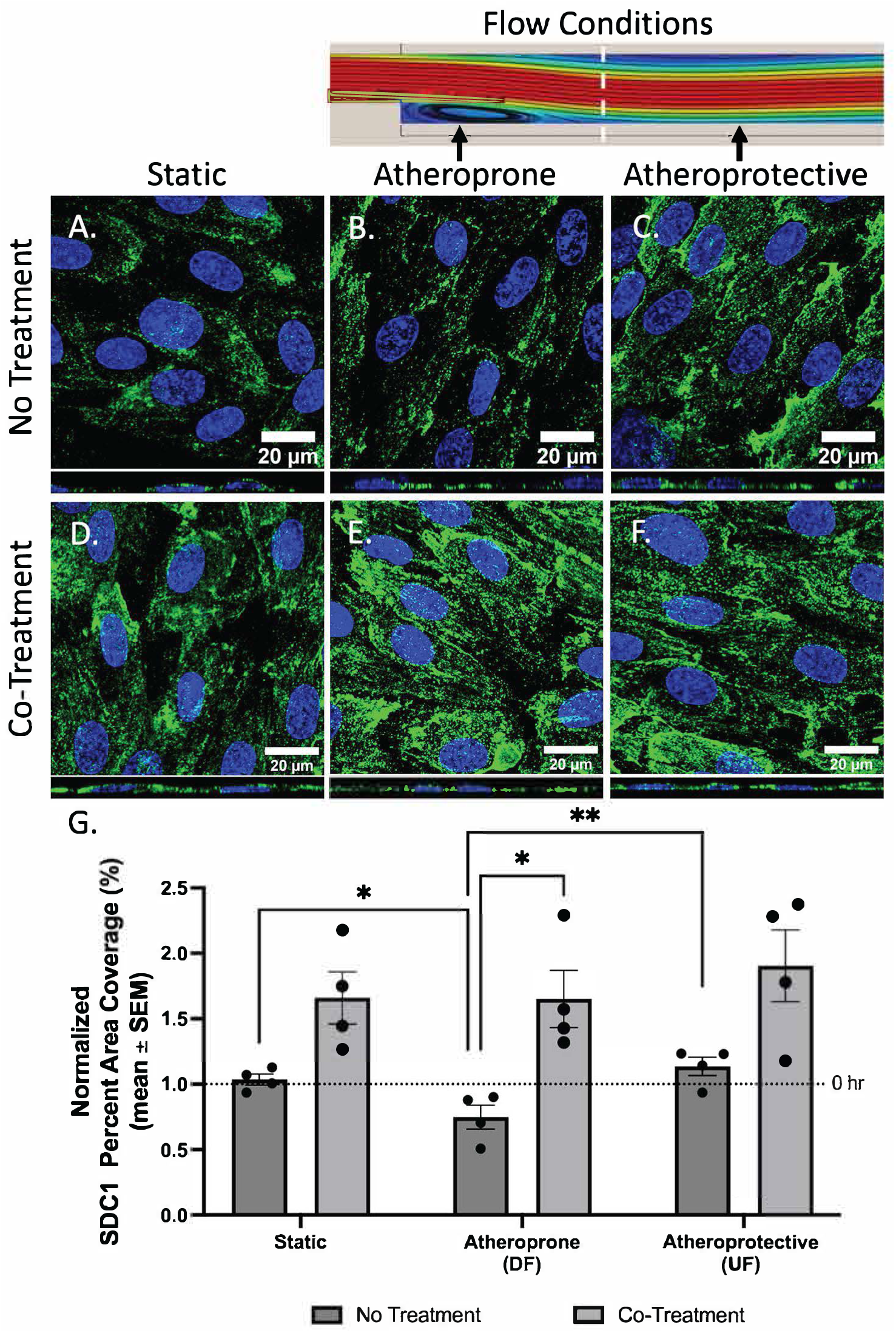
Effect of co-treatment on SDC1 (green), a GCX core protein, expression in HCAECs after 12 hours of flow at 12 dynes/cm^2^. Shown are 63x representative images of HCAECs in each experimental group. SDC1 expression increases in all conditions when the co-treatment is introduced to HCAECs. Blue represents cell nuclei labeled with DAPI. The scale bar is 20 μm. HCAECs in the no treatment group under A) static conditions, B) atheroprone conditions, and C) atheroprotective conditions. HCAECs in the co-treatment group under D) static conditions, E) atheroprone conditions, and F) atheroprotective conditions. G) Graph shows the mean ± SEM of percent area coverage of SDC1 staining normalized to the control (0 hr static condition; n = 4). Statistical analysis was performed using a two-way ANOVA. Significance is denoted by asterisks: *p<.05, **p<.01, ***p<.001, and ****p<.0001.

Figure 5 shows results from the *in vivo* partially ligated LCA murine model, including representative core protein SDC1 (red) stained images acquired at 40x magnification. The quantification that is shown reveals for ligated LCA, normalized to the RCA per condition, that SDC1 percent area fraction is 0.41 ± 0.04 in untreated (no vehicle) conditions, 0.55 ± 0.06 in vehicle-only conditions, and 0.84 ± 0.06 in co-treatment conditions, with co-treatment yielding statistically significantly better SDC1 expression compared to no treatment and vehicle-only. Analysis was also conducted on SDC1 transversal thickness of the ligated LCA normalized by non-ligated RCA for all mice observed. We report normalized SDC1 thicknesses of 0.72 ± 0.05 in untreated (no vehicle) conditions, 0.85 ± 0.05 in vehicle-only conditions, and most significantly significant 1.09 ± 0.05 in co-treatment conditions.

**Figure 5.**
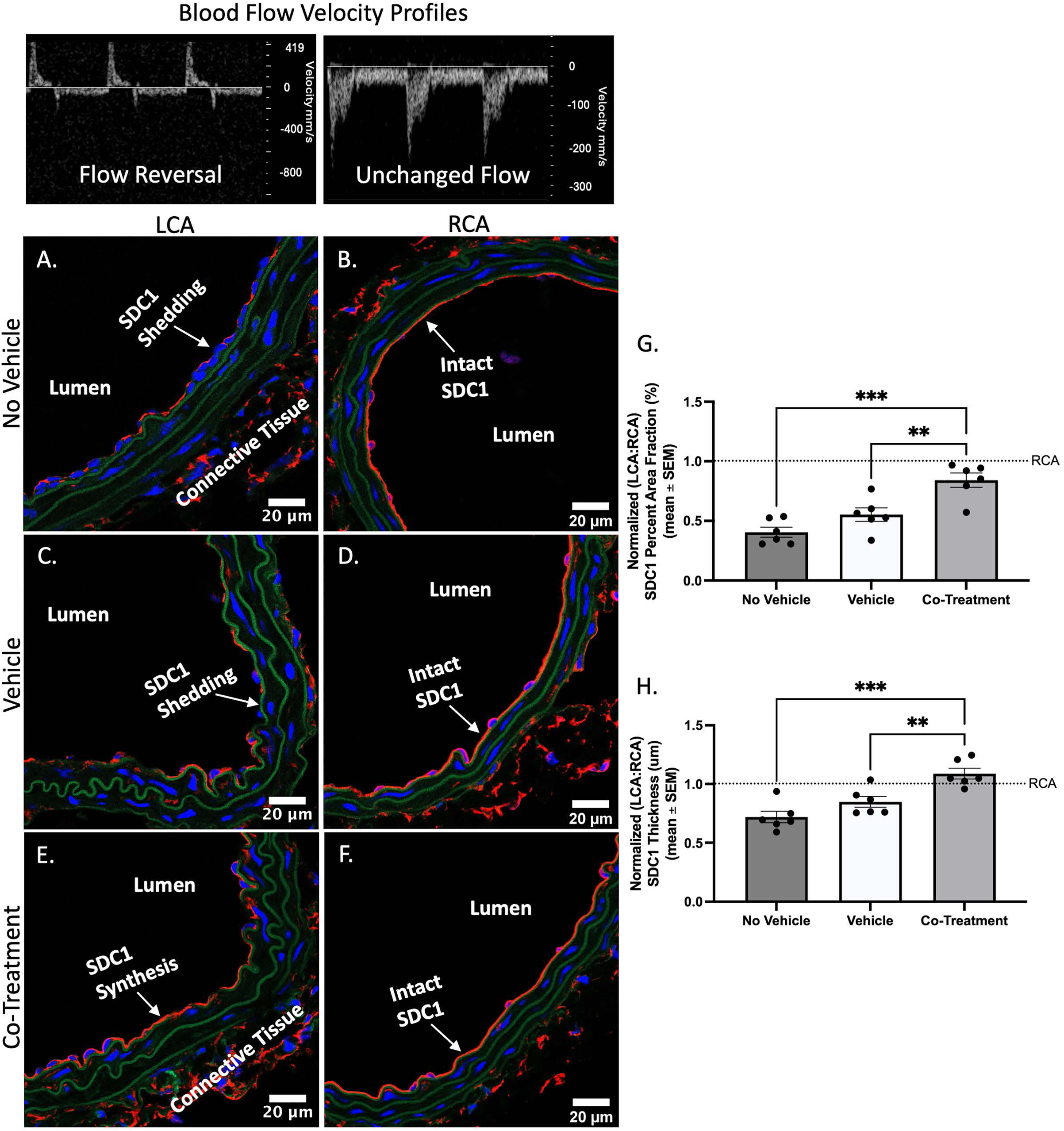
Effect of co-treatment on SDC1 (red) expression in murine carotid arteries. Shown are representative 40x images of carotid arteries from each cohort. The percent area coverage and thickness of SDC1 increased in the co-treatment group following a single dose of the co-treatment administered via retro-orbital injection. Blue represents cell nuclei labeled with DAPI. Green represents elastin. The scale bar is 20 μm. A) Ligated LCA from the no vehicle group. B) Control RCA from the no vehicle group. C) Ligated LCA from the vehicle only group. D) Control RCA from the vehicle only group. E) Ligated LCA from the co-treatment group. F) Control RCA from the co-treatment group. G) Graph shows the mean ± SEM of percent area coverage of SDC1 staining normalized to the control (RCA of each mouse; n = 6). H) Graph shows the mean ± SEM of SDC1 thickness normalized to the control (RCA of each mouse; n = 6). Statistical analysis was performed using a one-way ANOVA. Significance is denoted by asterisks: *p<.05, **p<.01, ***p<.001, and ****p<.0001.

Figure 6 shows *in vitro* results for GPC1, including both fluorescent (green for GPC1) images and plotted quantification of the 12-hour results normalized to 0-hour baseline conditions, as well as statistical analysis. Normalized GPC1 percent area coverage of untreated cultured HCAEC layers was found to be 1.08 ± 0.08 in 12-hour static conditions, 0.78 ± 0.08 in 12-hour DF conditions, and 1.25 ± 0.10 in 12-hour UF conditions. When co-treatment was introduced, the same cells were found to be covered by GPC1 at normalized levels of 1.29 ± 0.11 for static conditions, 1.32 ± 0.05 for DF conditions, and 1.35 ± 0.13 for UF conditions. With this, the only statistically significant co-treatment induced GPC1 enhancement occurred in DF conditions, shown by an approximate 70% increase.

**Figure 6.**
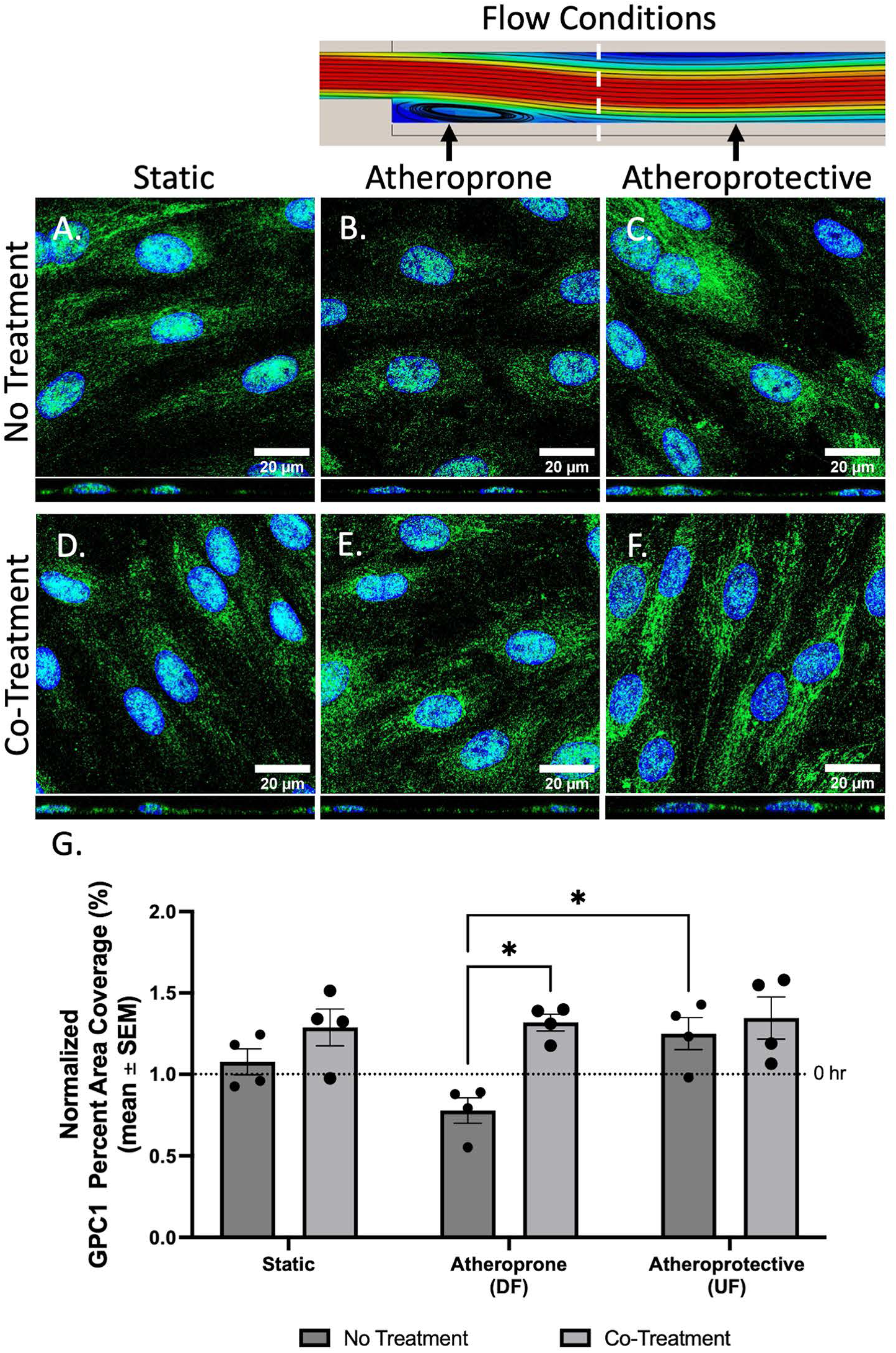
Effect of co-treatment on GPC1 (green), a GCX core protein, expression in HCAECs after 12 hours of flow at 12 dynes/cm^2^. Shown are representative 63x images of HCAECs in all experimental groups. The percent area coverage of GPC1 increased in atheroprone and atheroprotective conditions when co-treatment is introduced to the flow system. Blue represents cell nuclei labeled with DAPI. The scale bar is 20 μm. HCAECs in the no treatment group under A) static conditions, B) atheroprone conditions, and C) atheroprotective conditions. HCAECs in the co-treatment group under D) static conditions, E) atheroprone conditions, and F) atheroprotective conditions. G) Graph shows the mean ± SEM of percent area coverage of GPC1 staining normalized to the control (0 hr static condition; n = 4). Statistical analysis was performed using a two-way ANOVA. Significance is denoted by asterisks: *p<.05, **p<.01, ***p<.001, and ****p<.0001.

GPC1 data collected from the mouse model is shown in Figure 7, micrographically and quantitatively. When normalized to the RCA condition, the GPC1 percent area fraction of the inner LCA wall was found to be 0.45 ± 0.06 in untreated (no vehicle) conditions, 0.49 ± 0.04 in vehicle-only conditions, and 1.06 ± 0.07 in co-treatment conditions. Therefore, the co-treatment of albumin-S1P and heparin resulted in statistically significant restoration of GPC1. As for transversal LCA GPC1 thickness, normalized by RCA data, it measured in at 0.79 ± 0.03 in untreated (no vehicle) conditions, 0.80 ± 0.04 in vehicle-only conditions, and 1.10 ± 0.04 when mice were exposed to co-treatment of albumin-S1P and heparin. The co-treatment statistically significantly boosted GPC1 thickness compared to baseline conditions, to the extent of 43% compared to the untreated (no vehicle) case and 37% compared to the vehicle-only case.

**Figure 7.**
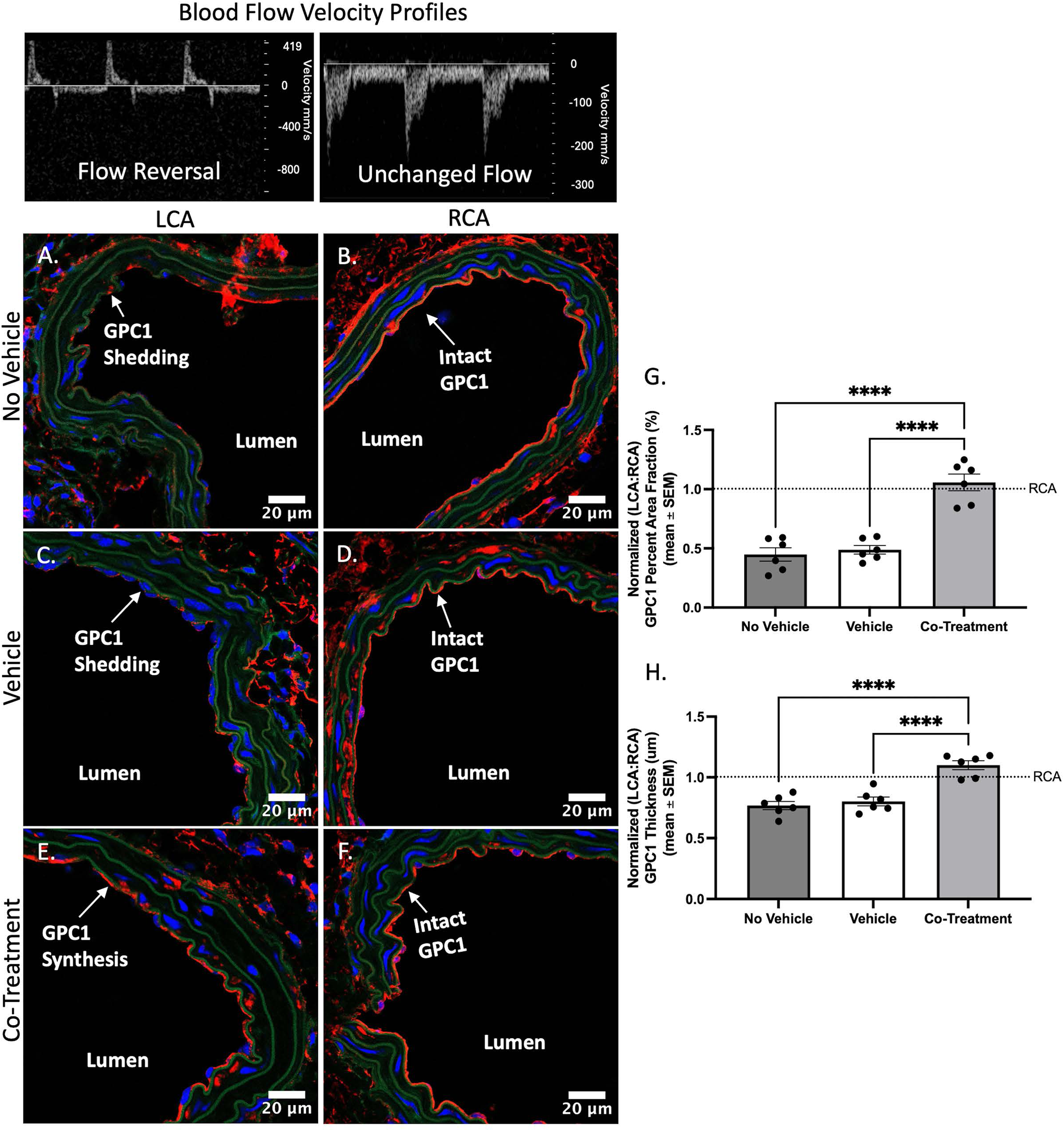
Effect of co-treatment on GPC1 (red) expression in murine carotid arteries. Shown are representative 40x images of carotid arteries of mice in all experimental cohorts. GPC1 percent area fraction and thickness increased in the co-treatment group following a single dose of the co-treatment administered via retro-orbital injection. Blue represents cell nuclei labeled with DAPI. Green represents elastin. The scale bar is 20 μm. A) Ligated LCA from the no vehicle group. B) Control RCA from the no vehicle group. C) Ligated LCA from the vehicle only group. D) Control RCA from the vehicle only group. E) Ligated LCA from the co-treatment group. F) Control RCA from the co-treatment group. G) Graph shows the mean ± SEM of percent area fraction of GPC1 staining normalized to the control (RCA of each mouse; n = 6). H) Graph shows the mean ± SEM of GPC1 thickness normalized to the control (RCA of each mouse; n = 6). Statistical analysis was performed using a one-way ANOVA. Significance is denoted by asterisks: *p<.05, **p<.01, ***p<.001, and ****p<.0001.

#### 3.2 Co-Treatment Impact on Endothelial and Vascular Function

##### Co-treatment of albumin-bound S1P and heparin does not affect flow-controlled in vitro cellular remodeling and morphology; cell alignment is not enhanced under any flow condition

Cell alignment analysis was conducted on brightfield images of HCAECs grown conditioned by static, DF, and UF conditions, with and without albumin-S1P and heparin co-treatment, as shown in Figure 8. The Figures 8G and 8H graphs report the OOP scores representing overall alignment of the HCAECs in the direction of flow and the MPA scores representing alignment of the HCAECs relative to each other. Recall the following, OOP = -1 means alignment perpendicular to the flow, OOP = +1 means alignment parallel to the flow, MPA = 1 indicates perfect alignment of all cells, and MPA = 0.5 indicates random orientations among cells. As seen in Figure 8, for both the OOP and MPA scores, the static flow condition showed the most random cell alignment, the DF condition showed a trend towards HCAEC alignment in the direction of flow and relative to each other, and the UF conditions showed the highest level of HCAEC alignment with flow and among HCAECs. Specifically, in the absence of treatment the OOP metrics, as shown in the box-and-whisker plots of Figure 8G, reveal that the static-conditioned HCAECs had a median of 0.17, with a range from -0.44 to 0.68 and a broad interquartile range (IQR) from -0.23 to 0.50. The DF-conditioned HCEACs had a higher median of 0.46, ranging from -0.41 to 0.83, and an IQR from 0.13 to 0.74. The UF-conditioned HCAECs exhibited the highest median at 0.70, with values from -0.04 to 0.91, and an IQR from 0.47 to 0.86. For static-conditioned HCAECs co-treated with albumin-S1P and heparin, compared to untreated static-conditioned HCAECs, we observed a lower median of -0.06 and a tighter range from -0.33 to 0.44, with an IQR from -0.27 to 0.15. The co-treatment did not seem to impact DF-conditioned HCAECs as seen by the maintained median of 0.47, range from -0.17 to 0.73 and an IQR from 0.09 to 0.61. Compared to untreated UF-conditions, the co-treated UF-conditioned group of HCAEC’s median slightly increased to 0.73, with values between -0.24 and 0.93, and an IQR from 0.38 to 0.88. Furthermore, in the absence of treatment the MPA metric was found to be 0.70 ± 0.02 in static conditions, 0.77 ± 0.01 for DF conditions, and 0.83 ± 0.02 for UF conditions. With albumin-S1P and heparin co-treatment the MPA metric was found to be 0.72 ± 0.003 in static conditions, 0.77 ± 0.01 in DF conditions, and 0.82 ± 0.02 in UF conditions. Neither the OOP nor MPA metric revealed a statistically significant improvement in cell alignment due to albumin-S1P and heparin co-treatment, for any of the flow conditions. These results indicate that, at least in the 12-hour time frame for this investigation, local cell elongation and alignment are primarily influenced by the flow condition, not the albumin-S1P and heparin co-treatment.

**Figure 8.**
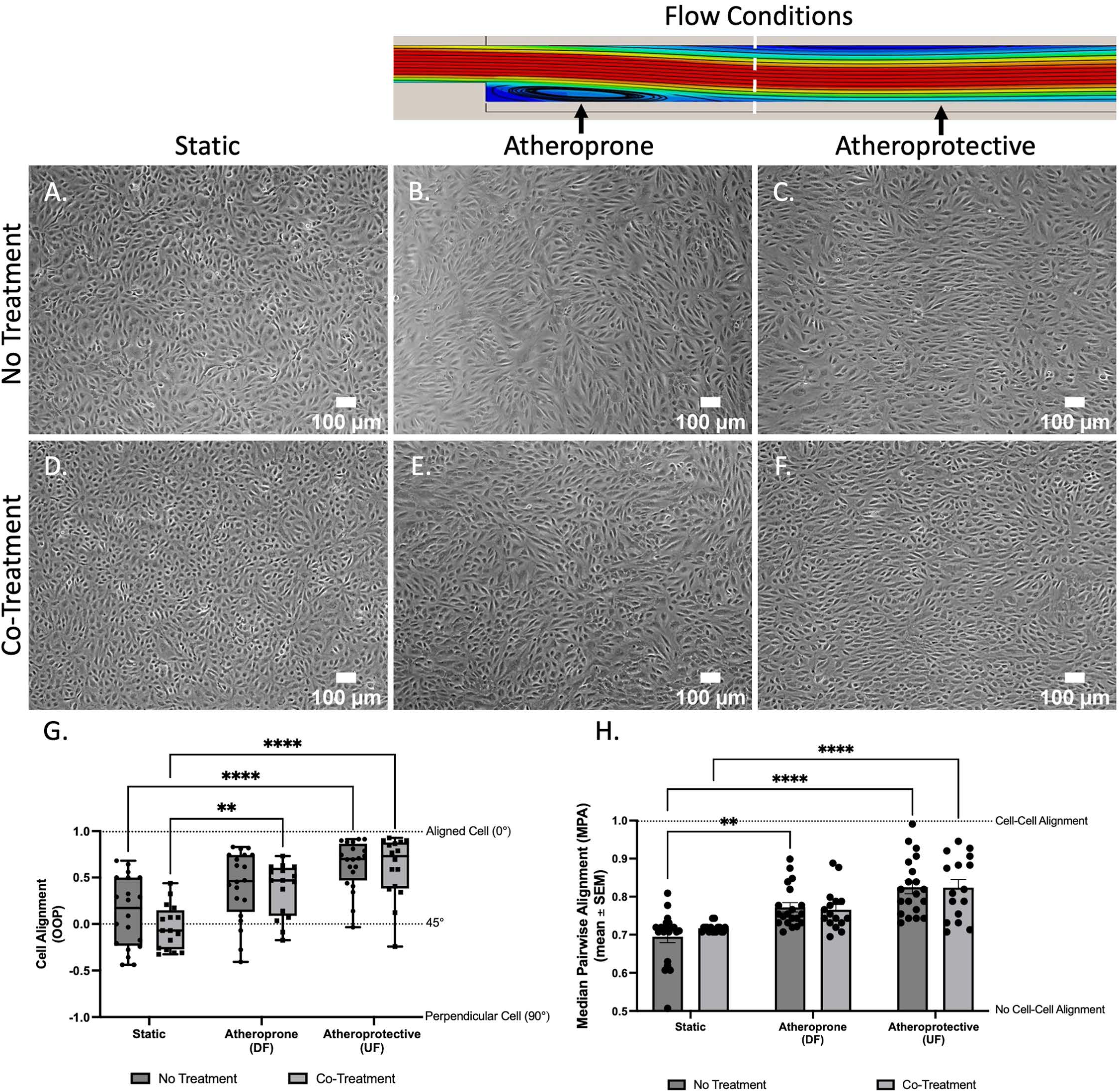
*In* vitro cellular remodeling as indicated by EC morphology via cell alignment in relation to flow. Representative phase contrast images of HCAECs at 10x are shown for all cohorts, which include HCAECs exposed to: A) static condition with no treatment, B) atheroprone region with no treatment, C) atheroprotective region with no treatment, D) static condition with co-treatment, (E) atheroprone region with co-treatment, and (F) atheroprotective region with co-treatment. All static and dynamic conditions (12 dynes/cm^2^) were for 12 hours. Cell alignment was analyzed in two ways: (G) Object Orientation Parameter (OOP), which determines cell alignment in relation to flow and (H) Median Pairwise Alignment (MPA), which determines cell alignment in relation to other ECs. Each data point on the graph represents the mean individual OOP or MPA from one flow experiment, in which each cohort as range of n = 16 – 20. Box and whisker plot in (G) for OOP shows the minimum, first quartile, median, third quartile, and maximum values for each condition. MPA in (H) shows the mean and error bars which represents the SEM. Statistical analysis was performed using a two-way ANOVA. Significance is denoted by asterisks: *p<.05, **p<.01, ***p<.001, and ****p<.0001.

##### Co-Treatment of albumin bound S1P and heparin improves vascular tone via upregulation of p-eNOS and reduction in vessel wall thickness

Vascular tone was studied *in vitro* by investigating p-eNOS and eNOS. Figure 9 shows representative images of the percent area fraction of p-eNOS (active eNOS; green) in HCAECs subject to different conditions for 12 hours, while Figure 10 shows representative images of total eNOS. The figures also show plots derived from analyzing the full set of image data and normalizing the data with respect to 0-hour static baseline conditions. As seen in Figure 9, in untreated conditions, normalized p-eNOS expression in static HCAECs was 1.53 ± 0.09, DF HCAECs was 1.00 ± 0.10, and UF HCAECs was 2.29 ± 0.28. Figure 10 shows normalized total eNOS expression in static HCAECs was 1.38 ± 0.36, dropped significantly to 0.705 ± 0.307 for DF HCAECs, and for UF HCAECs was 1.54 ± 0.457. These p-eNOS and eNOS trends are to be expected, based on the well-understood relative effects of various flow conditions on p-eNOS and eNOS [11,16,50,51]. Upon exposure to co-treatment with albumin-S1P and heparin, p-eNOS expression by static HCAECs rose to 1.69 ± 0.31, DF HCAECs rose to 2.18 ± 0.45, and UF HCAECs remained nearly consistent at 2.00 ± 0.16 (Figure 9). Clearly, the impact of the co-treatment on p-eNOS is the most statistically significant in DF conditions (Figure 9). Interestingly, no statistical significance was observed in HCAECs in any flow condition (static, DF, or UF) when comparing untreated conditions to co-treatment conditions (Figure 10). This signifies that the albumin-S1P and heparin co-treatment does not impact total eNOS expression in HCAECs; it primarily exerts its impact on p-eNOS.

**Figure 9.**
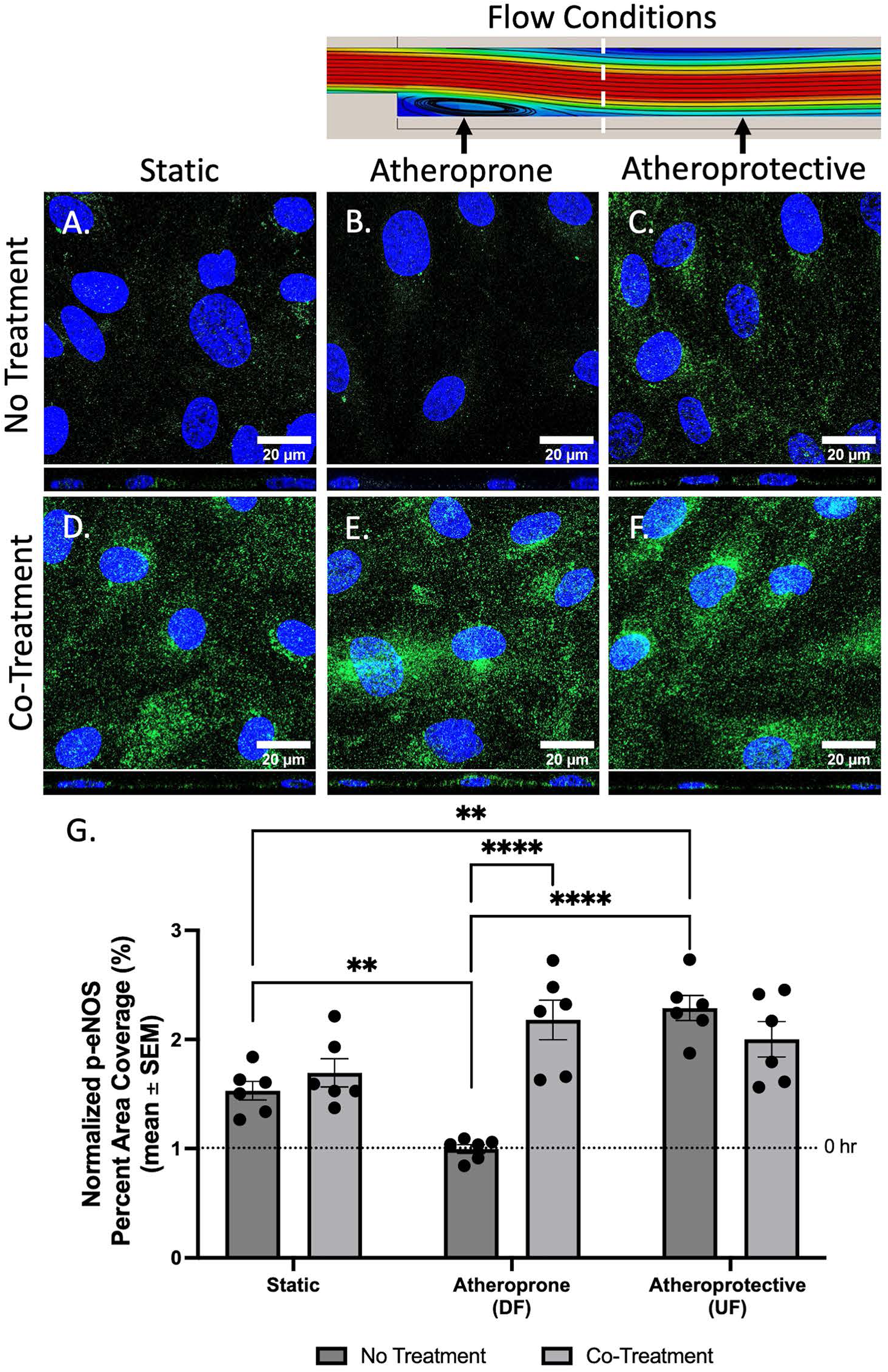
Effect of co-treatment on activated eNOS (p-eNOS; green) expression in HCAECs after 12 hours of flow at 12 dynes/cm^2^. Shown are representative 63x images of HCAECs from each experimental group. The percent area coverage of p-eNOS increased when the co-treatment is administered to the flow system. Blue represents cell nuclei labeled with DAPI. The scale bar is 20 μm. HCAECs in the no treatment group under A) static conditions, B) atheroprone conditions, and C) atheroprotective conditions. HCAECs in the co-treatment group under D) static conditions, E) atheroprone conditions, and F) atheroprotective conditions. G) Graph shows the mean ± SEM of percent area coverage of p-eNOS staining normalized to the control (0 hr static condition; n = 6). Statistical analysis was performed using a two-way ANOVA. Significance is denoted by asterisks: *p<.05, **p<.01, ***p<.001, and ****p<.0001. *Note: Supplemental Figure 4 contains results that show p-eNOS expression in HCAECs that were treated with either albumin-bound S1P or heparin separately*.

**Figure 10.**
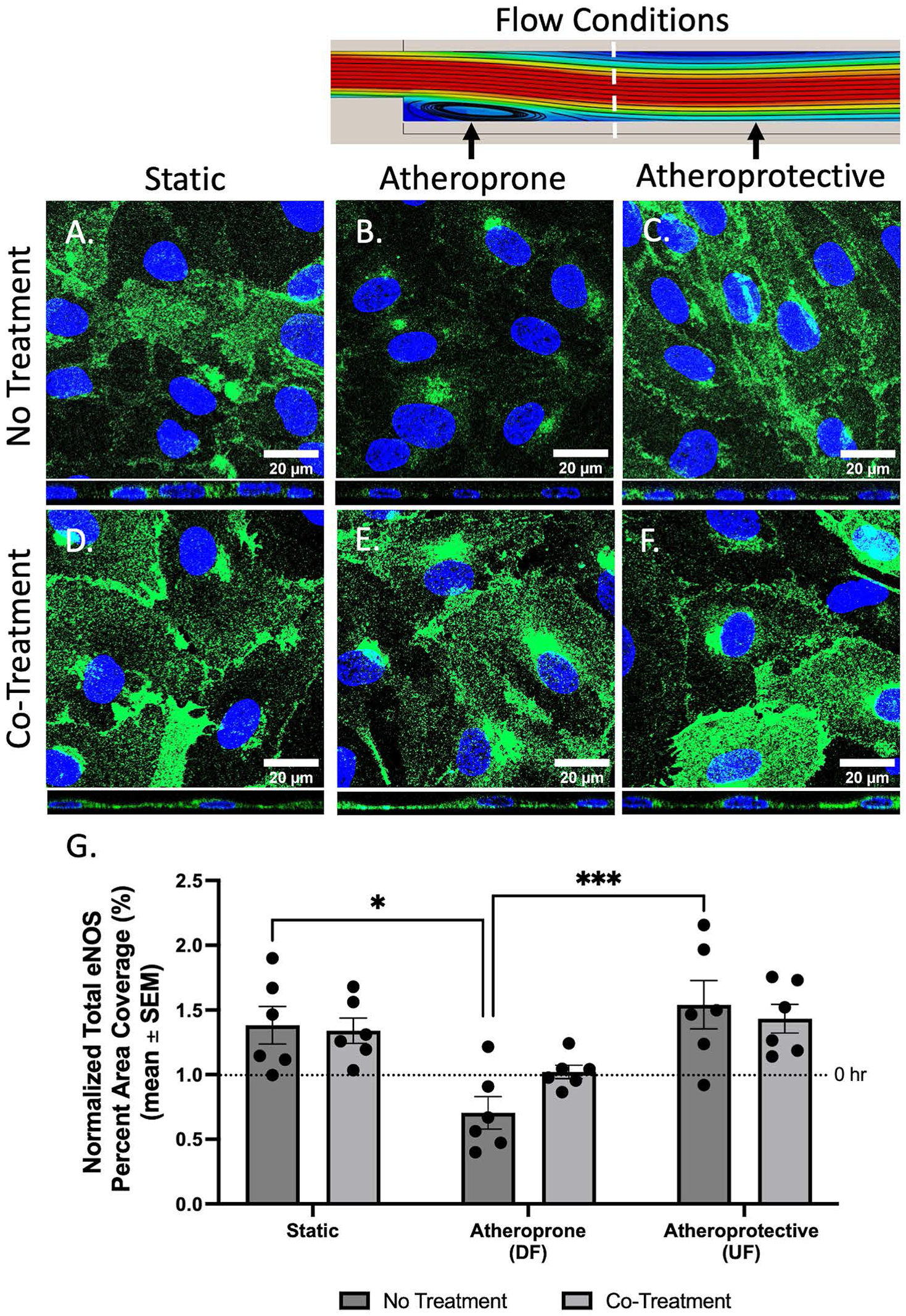
Effects of co-treatment on total eNOS (green) expression in HCAECs after 12 hours of flow at 12 dynes/cm^2^. Shown are 63x representative images of total eNOS in HCAECs in different experimental groups. No changes were present in the percent area coverage of total eNOS with or without co-treatment. Blue represents cell nuclei labeled with DAPI. The scale bar is 20 μm. HCAECs in the no treatment group under A) static conditions, B) atheroprone conditions, and C) atheroprotective conditions. HCAECs in the co-treatment group under D) static conditions, E) atheroprone conditions, and F) atheroprotective conditions. G) Graph shows the mean ± SEM of percent area coverage of total eNOS staining normalized to the control (0 hr static condition; n = 6). Statistical analysis was performed using a two-way ANOVA. Significance is denoted by asterisks: *p<.05, **p<.01, ***p<.001, and ****p<.0001. *Note: Supplemental Figure 5 contains results that show total eNOS expression in HCAECs that were treated with either albumin-bound S1P or heparin separately*.

To study vascular tone *in vivo*, vessel remodeling was investigated by periodically observing vessel wall thickness (mm) and systolic and diastolic vessel diameters (mm) via ultrasound. Figure 11 presents data collected five days post-ligation of the left carotid artery (LCA) of mice, with the right carotid artery (RCA) left intact to serve as a reference. Focusing on LCA wall thickness (Figure 11B), the untreated LCA wall thickness was 0.07 ± 0.01 mm, while the RCA wall thickness was 0.05 ± 0.003 mm. For the vehicle-only group, the LCA wall was 0.06 ± 0.002 mm thick compared to 0.04 ± 0.002 mm for the RCA. In the albumin-S1P and heparin co-treated group, the LCA wall measured 0.05 ± 0.004 mm, with the RCA at 0.05 ± 0.002 mm. These results indicate that the LCA wall is significantly thicker than the RCA wall in both untreated and vehicle-only conditions, but co-treatment renders LCA thickness statistically similar to RCA thickness. Figures 11C and 11D show the systolic and diastolic vessel diameters in mice across various experimental cohorts. The systolic vessel diameter of the ligated LCA was 0.53 ± 0.02 mm in untreated mice, 0.48 ± 0.01 mm in vehicle-only mice, and 0.48 ± 0.01 mm in co-treated mice (Figure 11C). The diastolic vessel diameter was 0.46 ± 0.02 mm for untreated mice, 0.42 ± 0.01 mm for vehicle-only mice, and 0.43 ± 0.02 mm for co-treated mice (Figure 11D). In the RCA, the systolic vessel diameter was 0.49 ± 0.02 mm in untreated conditions, 0.53 ± 0.02 mm in vehicle-only conditions, and 0.50 ± 0.02 mm in co-treated conditions (Figure 11C). The RCA diastolic vessel diameter was 0.40 ± 0.02 mm in untreated conditions, 0.46 ± 0.02 mm in vehicle-only conditions, and 0.45 ± 0.02 mm in co-treated conditions (Figure 11D). No statistically significant differences were observed across all experimental conditions for systolic and diastolic diameters. Since the systolic and diastolic diameters of the LCA and RCA remained similar across all experimental conditions, no discernible impact of co-treatment on vessel diameters was observed. This indicates that the co-treatments impact on vascular tone is most effectively exerted by normalizing vascular wall thickness.

**Figure 11.**
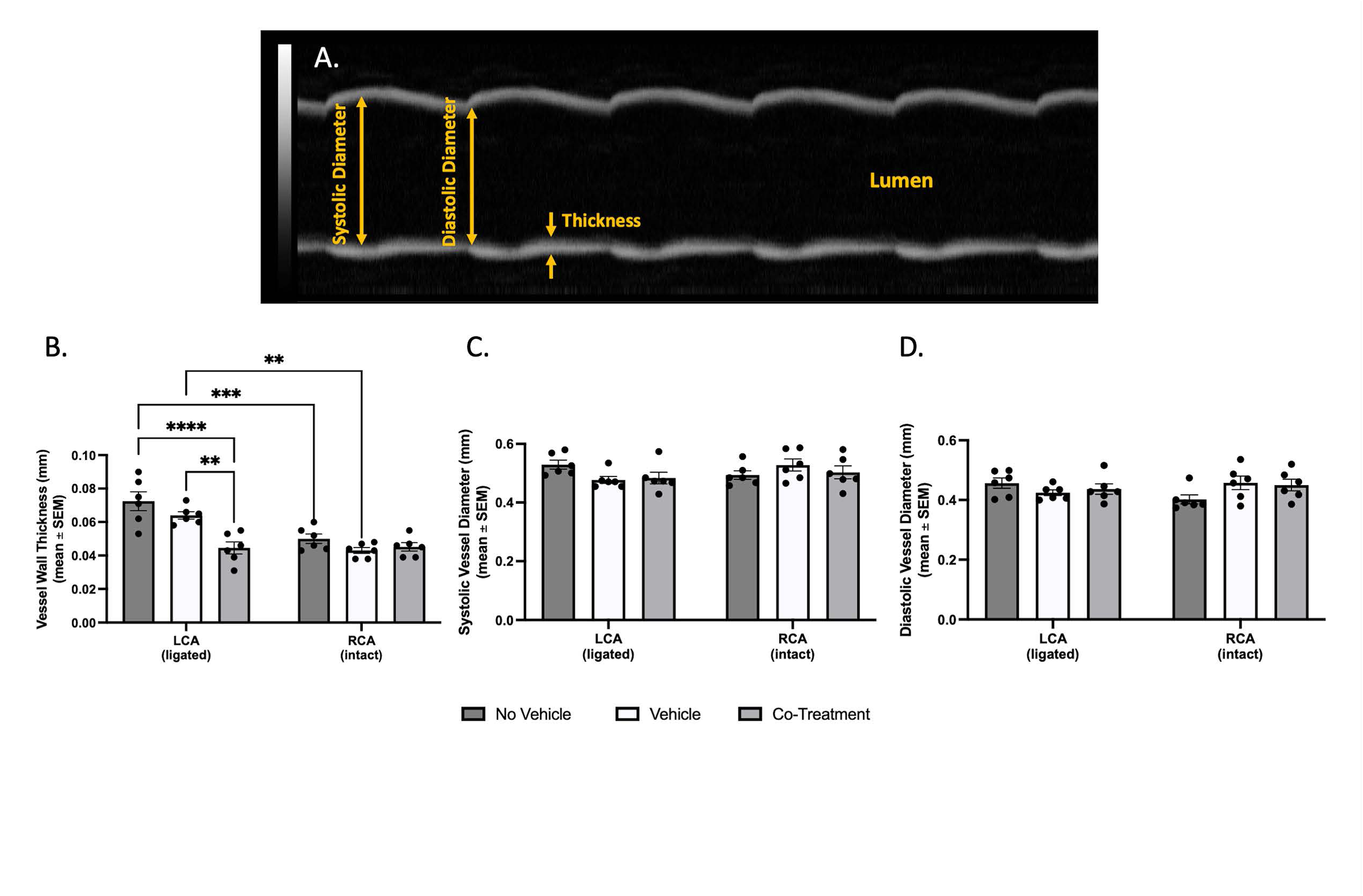
Effect of co-treatment on vascular tone as indicated by (B) vessel wall thickness (mm), (C) systolic vessel diameter (mm), and (D) diastolic vessel diameter (mm) via ultrasound images *in vivo* for experimental cohorts. Co-treatment was able to provide vessel remodeling after a single retro-orbital IV injection in mice shown by reduction in vessel wall thickness; however, other parameters were not impacted. (A) Provides a representative image of a carotid artery in B-mode in longitudinal axis and how different parameters were collected. These data points were collected 5 days-post ligation of LCA and mice were either given no vehicle, vehicle only, or co-treatment. Each data point on the graphs represents the mean individual for mouse in each cohort, where each cohort has an n = 6. The mean and error bars representing the SEM. Statistical analysis was performed using a two-way ANOVA. Significance is denoted by asterisks: *p<.05, **p<.01, ***p<.001, and ****p<.0001.

##### Co-Treatment of albumin bound S1P and heparin reduces inflammation and vessel hyperpermeability in atheroprone region in vivo

Figure 12 shows 40x representative images of positive-CD68 immunofluorescence, which identified macrophages taken up into the vessel walls of the carotid arteries in all cohorts of mice. A percent area fraction was calculated to determine macrophage infiltration, specifically within vessel walls of carotid arteries.

**Figure 12.**
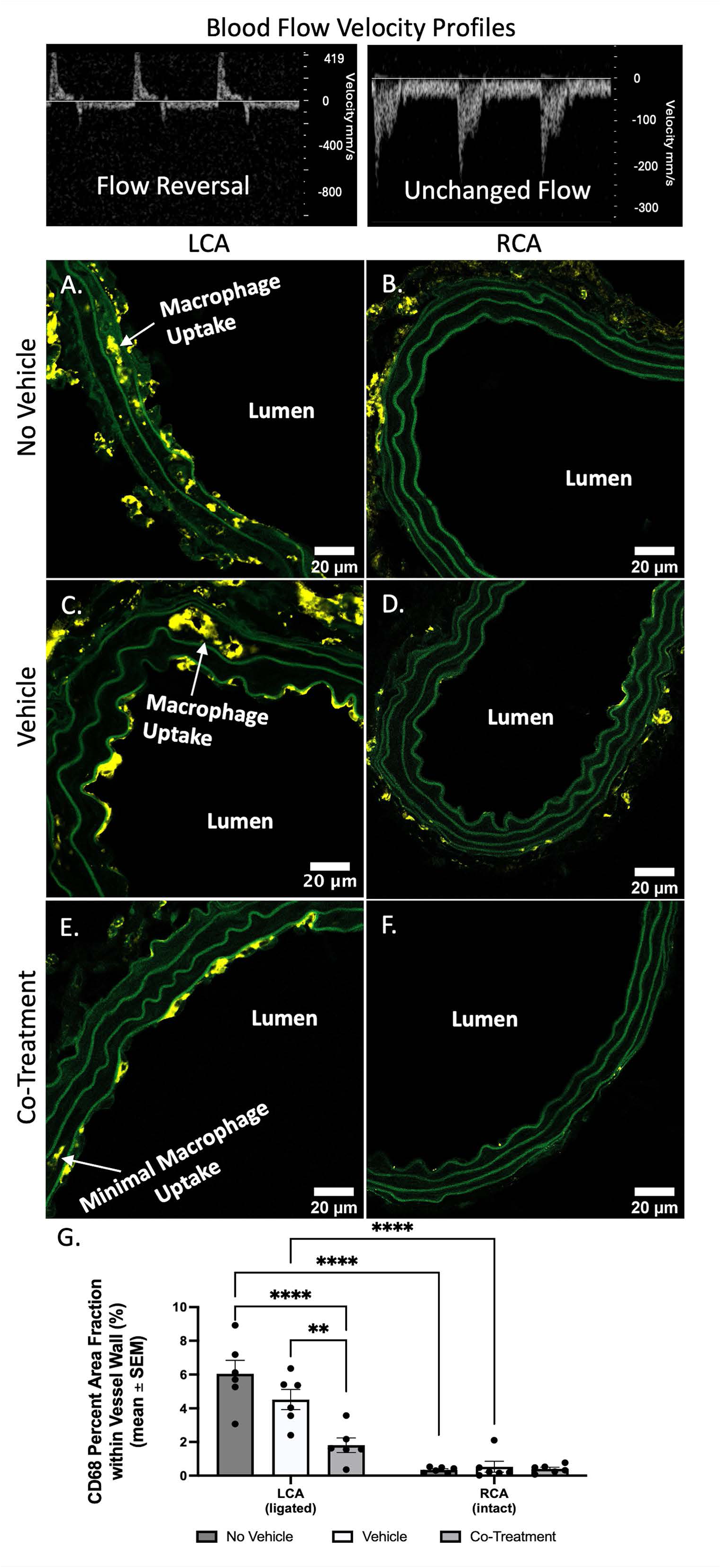
Effect of co-treatment on CD68 expression in murine carotid arteries. Shown are 40x representative carotid arteries of mice in different experimental cohorts. Staining of CD68 (yellow), a protein expressed highly by circulating macrophages, was used to evaluate barrier permeability and macrophage uptake into the carotid arteries. CD68 uptake within vessel walls decreased in the co-treatment group following a single dose of the co-treatment administered via retro-orbital injection. Green represents elastin. The scale bar is 20 μm. A) Ligated LCA from the no vehicle group. B) Control RCA from the no vehicle group. C) Ligated LCA arterial ring from the vehicle only group. D) Control RCA arterial ring from the vehicle only group. E) Ligated LCA from the co-treatment group. F) Control RCA from the co-treatment group. G) Graph shows the mean ± SEM of percent area fraction of CD68 staining within vessels walls of the LCA and RCA (n = 6). Statistical analysis was performed using a two-way ANOVA. Significance is denoted by asterisks: *p<.05, **p=.01, ***p<.001, and ****p<.0001.

Macrophage uptake in the ligated LCA of untreated (no vehicle) mice was 6.05 ± 0.80%, vehicle-only mice was 4.52 ± 0.60%, and mice administered the co-treatment was 1.81 ± 0.43%. These outcomes were found to be statistically significantly different. In contrast to the ligated LCA vessels, there was far less macrophage uptake in the non-ligated RCA vessels. RCA vessels were infiltrated by macrophages to the extent of 0.35 ± 0.07% in untreated (no vehicle) conditions, 0.53 ± 0.32% in vehicle-only conditions, and 0.40 ± 0.10% in co-treatment conditions. No statistical differences were observed across all cohorts when observing macrophage infiltration of the reference RCA in mice.

#### 3.3 Co-Treatment Mechanism of Action

##### Co-treatment of albumin-bound S1P and heparin circulates longer and at higher concentrations in regions of DF

Computational fluid simulations confirm a region of recirculating flow downstream of the step within the *in vitro* parallel-plate flow chamber device (Figure 13). To investigate the impact of the step on particle residence time, particles were uniformly released throughout the domain illustrated by the black rectangle in Figure 13 and counted the number of particles still suspended within the fluid at *t* = 0.2 *s* after being released. It was determined that the particle concentration within the circulating region was notably greater than that outside of the recirculating region (Figure 13c). The greater concentration in the recirculating region is indicative that particles in this domain tend to reside longer than particles outside of this region, which are downstream of the computational domain illustrated. This corroborates the notion that the co-treatment has a higher concentration and longer residence time in the atheroprone DF region *in vitro*.

**Figure 13.**
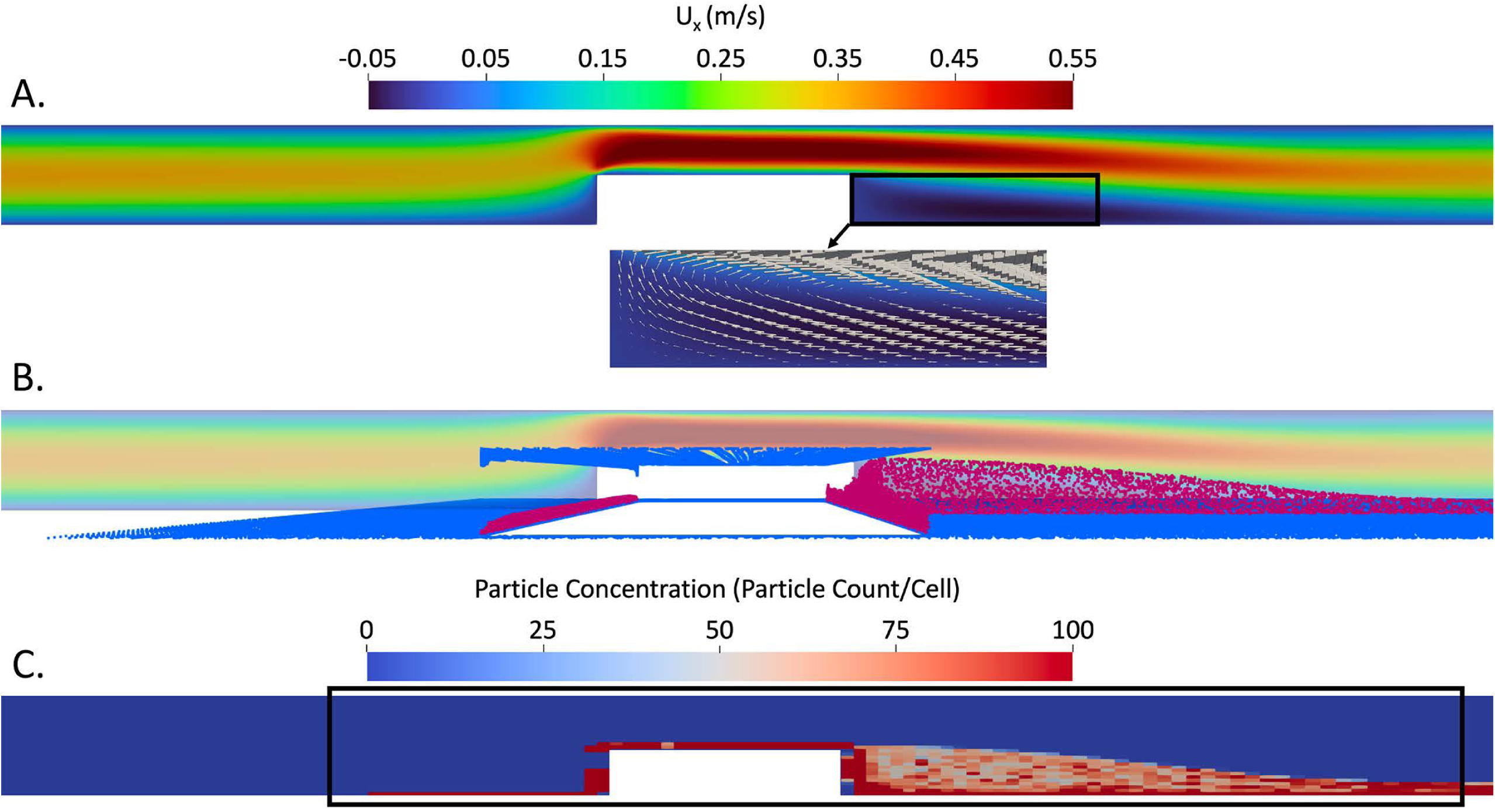
Fluid and particle dynamic simulations were conducted on parallel-plate flow chamber with co-treatment circulating the *in vitro* apparatus. (A) Computational flow field around the step in the microfluidic device. Simulations show a clear region of recirculating flow downstream of the step. Black arrows illustrate the velocity vector field in the recirculating region. (B) Computational particle analysis of co-treatment (red particles) circulating parallel-plate flow chamber. (C) Particle transport simulations demonstrate that particles released within the recirculating flow region reside there for longer periods of time, causing a greater particle concentration relative to particles released outside of the recirculating region (snapshot from t=0.2 s following release). The black rectangle indicates the regions where particles were released. Note that a coarser mesh is used for visualization purposes only.

##### Co-treatment of albumin-bound S1P and heparin appears to restore S1PR1 expression in regions of DF

As previously mentioned, S1PR1 is critical for EC mechanotransduction, regulation of endothelial barrier maintenance, vascular tone, and inflammation [34,52]. S1PR1-associated signaling pathways may explain the mechanism by which albumin-S1P and heparin co-treatment restores endothelial barrier integrity and vascular tone in atheroprone-DF region. Figure 14 presents representative images of S1PR1 expression (green) in HCAECs exposed for 12 hours to different mechanical stimuli, with or without treatment. The data points in Figure 14G are normalized with respect to baseline conditions at 0 hours. Without treatment, normalized S1PR1 expression was found to be 1.14 ± 0.16 under static conditions, 0.80 ± 0.05 under DF conditions, and 1.62 ± 0.12 under UF conditions. The difference between DF and UF S1PR1 expression is statistically significant. With albumin-S1P and heparin co-treatment, compared to untreated results, HCAEC S1PR1 expression increased significantly to 2.42 ± 0.352 under static conditions, increased to 1.30 ± 0.19 under DF conditions, and remained sustained with 1.90 ± 0.28 under UF conditions. These findings suggest that S1PR1 expression can be enhanced in DF regions through co-treatment, potentially activating S1PR1-associated signaling pathways to improve endothelial health.

**Figure 14.**
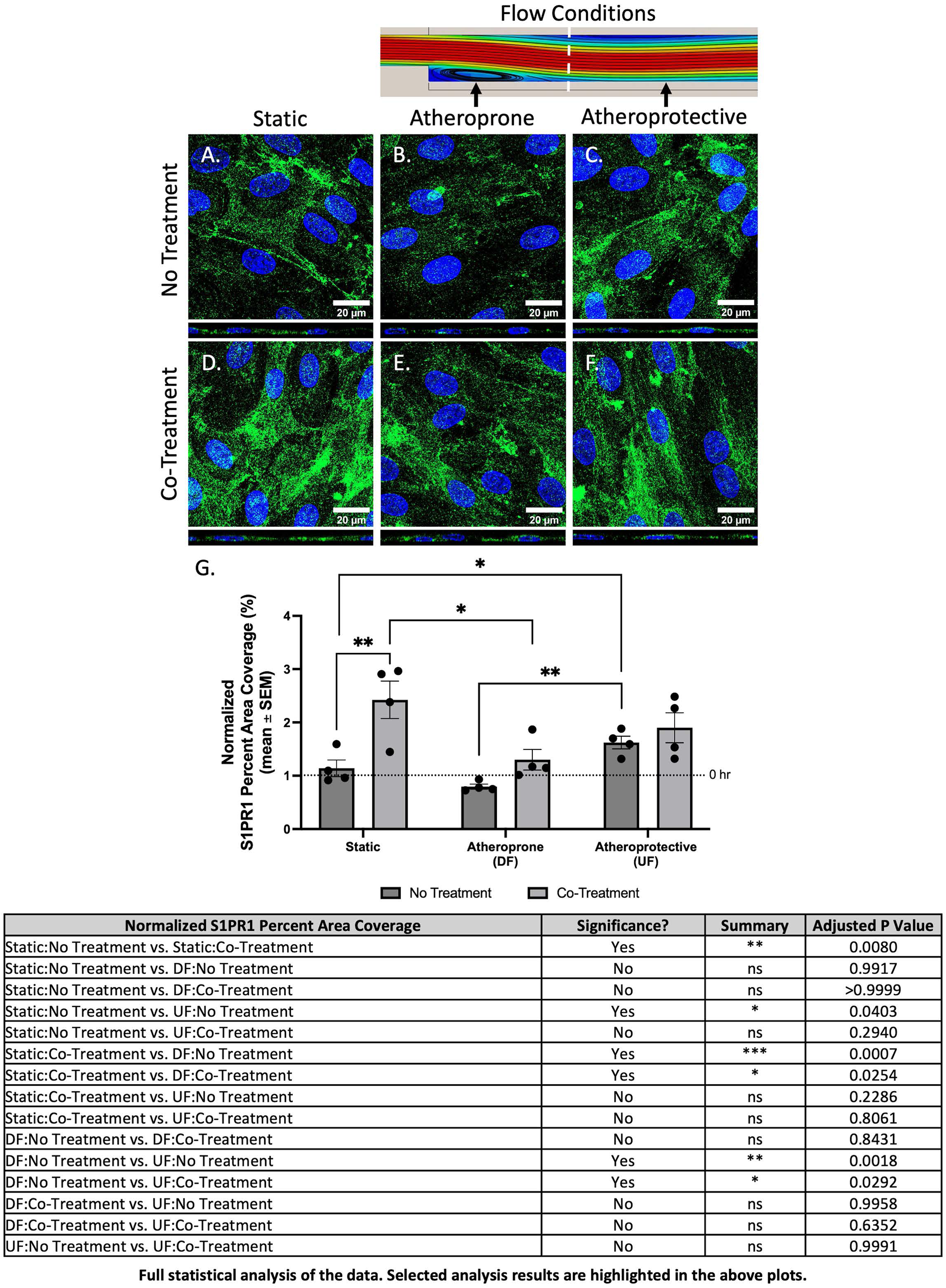
Effect of co-treatment on S1PR1 (green) expression in HCAECs after 12 hours of flow at 12 dynes/cm^2^. Shown are representative 63x images of S1PR1 expression in HCAECs amongst different experimental groups. S1PR1, a G-protein coupled receptor found specifically on ECs, was used to determine its involvement in the therapeutic mechanism of action, and its expression increased in the atheroprotective and atheroprone regions when the co-treatment is introduced to the flow system, signifying activation. Blue represents cell nuclei labeled with DAPI. The scale bar is 20 μm. HCAECs in the no treatment group under A) static conditions, B) atheroprone conditions, and C) atheroprotective conditions. HCAECs in the co-treatment group under D) static conditions, E) atheroprone conditions, and F) atheroprotective conditions. G) Graph shows the mean ± SEM of percent area coverage of S1PR1 staining normalized to the control (0 hr static condition; n = 4). Statistical analysis was performed using a two-way ANOVA. Significance is denoted by asterisks: *p<.05, **p<.01, ***p<.001, and ****p<.0001.

##### Co-treatment of albumin-bound S1P and heparin upregulates expression of EXTL3

*EXTL3*, as previously mentioned, is part of the exostosin family of genes that encodes glycosyltransferases involved in HS biosynthesis [53]. *EXTL3* has dual functions: it adds N-acetylglucosamine (GlcNAc) to the protein linkage region of GAGs, which is its GlcNAc-TI activity, and to the growing HS chain, which is its GlcNAc-TII activity [53]. This makes *EXTL3* an enzyme that plays a crucial role in both the initiation and elongation of HS chains [53]. Upregulation of *EXTL3* after co-treatment could potentially synthesize new endothelial glycocalyx (GCX) in atheroprone disturbed flow (DF) regions. Figure 15 shows 40x images of *EXTL3* (red) in the endothelium of carotid arteries across all mouse cohorts. The percentage area fraction of *EXTL3* on the carotid artery endothelium was calculated: in the partially ligated LCA, it was 7.90 ± 0.640% with no treatment, 7.72 ± 1.04% with vehicle, and 13.1 ± 1.46% with co-treatment. In the non-ligated RCA it was 17.0 ± 1.38% with no treatment, 16.4 ± 1.34% with vehicle, and 13.2 ± 1.08% with co-treatment. The RCA served as the control for each mouse, normalizing LCA data to assess fold change in *EXTL3* expression. Normalized *EXTL3* expression in LCA vessels, relative to RCA, was 0.48 ± 0.06 in untreated conditions, 0.46 ± 0.03 in vehicle-only conditions, and 0.98 ± 0.06 in co-treatment conditions. The increase in *EXTL3* expression with co-treatment was statistically significant compared to both untreated and vehicle-only conditions.

**Figure 15.**
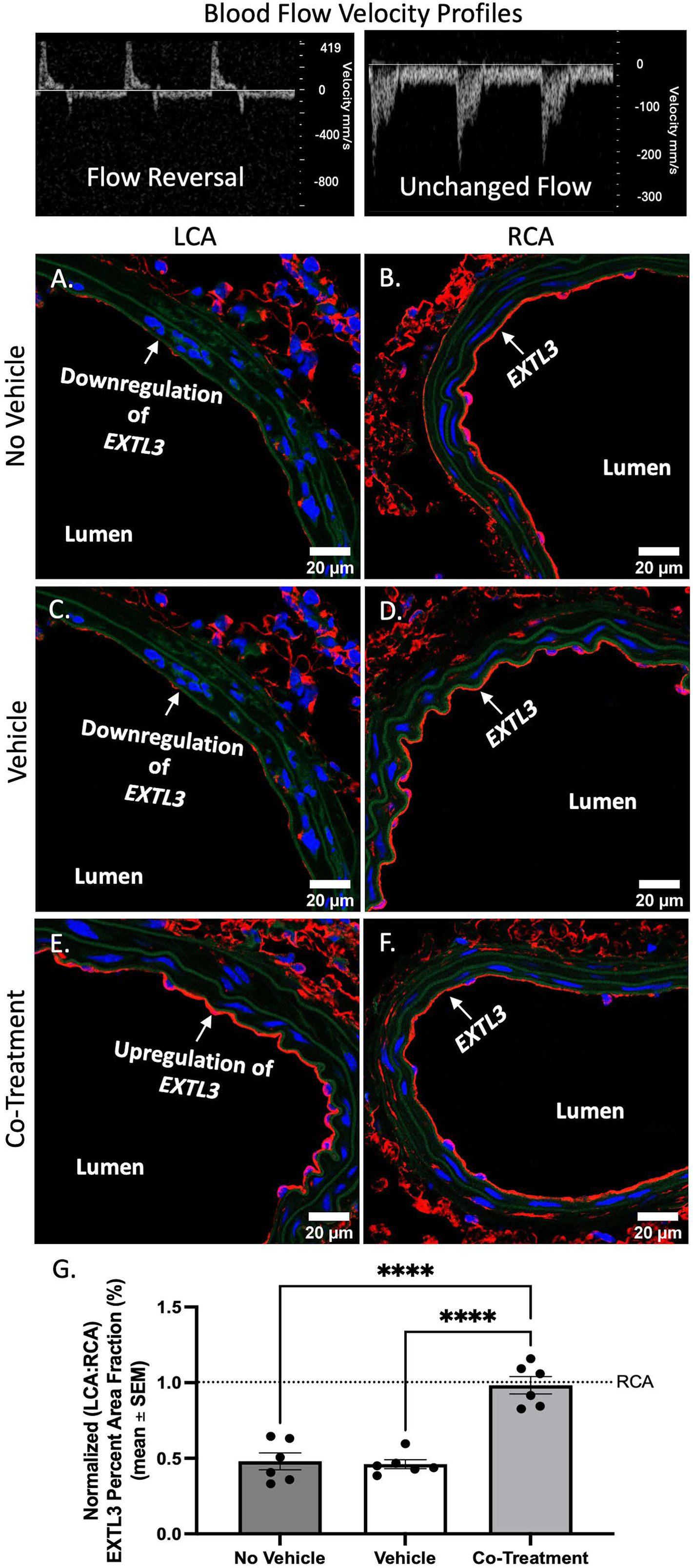
Effect of the co-treatment on *EXTL3* (red) expression in murine carotid arteries. Shown are 40x representative images of carotid arteries from mice from all experimental cohorts. *EXTL3* (red), which encodes the glycosyltransferases responsible for the biosynthesis of glycosaminoglycan heparan sulfate, was used to explore a potential mechanism of action for the co-treatment. The percent area fraction of *EXTL3* increases in the co-treatment group following a single dose of co-treatment administered via retro-orbital injection, signifying its activation. Green represents elastin. Blue represents cell nuclei. The scale bar is 20 μm. A) Ligated LCA from the no vehicle group. B) Control RCA from the no vehicle group. C) Ligated LCA from the vehicle only group. D) Control RCA from the vehicle only group. E) Ligated LCA from the co-treatment group. F) Control RCA from the co-treatment group. G) Graph shows the mean ± SEM of percent area fraction of *EXTL3* staining normalized to the control (RCA of each mouse). The mean and error bars representing the SEM are included for each group with n=6. Statistical analysis was performed using a two-way ANOVA. Significance is denoted by asterisks: *p<.05, **p=.01, ***p<.001, and ****p<.0001.

### 4. DISCUSSION

This study showed that treating cells with albumin-bound S1P and heparin together can restore the integrity of the GCX and improve endothelial function in regions of the vasculature prone to atherosclerosis. Evidence of GCX improvement came from WGA-labeled GCX, HS, and core proteins like SDC1 and GPC1. Restored endothelial function was marked by improvements in vascular tone and barrier functionality. While previous research has shown that S1P and heparin separately improve EC GCX integrity [36,54–57], our study is the first to combine these components and demonstrate their effects in complex and clinically relevant models. Specifically, we found that the co-treatment enhances GCX integrity in primary human coronary artery ECs (HCAECs), which are highly relevant to cardiovascular disease. Additionally, in vivo application of the co-treatment in a ligated left carotid artery (LCA) mouse model under DF conditions showed significant improvements in GCX integrity. Mechanistically, our study is the first to reveal that the co-treatment upregulates EXTL3, the key enzyme involved in HS synthesis *in vivo*.

We believe the albumin-bound S1P and heparin co-treatment has pleiotropic effects on EC GCX health by synthesizing and protecting GCX structure. For instance, Zeng et al. [36,56] have shown that S1P prevents SDC1 shedding via inhibition of activation of matrix metalloproteinases and synthesizes the GCX via phosphoinositide 3-kinase (PI3K) pathway, particularly in rat fat pad ECs. Additionally, heparin is postulated to protect the EC GCX from degradation, particularly in sepsis, by acting as an inhibitor of heparanase, an enzyme that specifically cleaves HS from the EC GCX [58]. This was confirmed in a preclinical study that reported GCX shedding in lung microvessels due to HS degradation induced by tumor necrosis factor-alpha (TNF-α) dependent heparanase activation [54]. This was attenuated by heparin treatment in a lipopolysaccharide murine model [54]. Furthermore, Potje et al. [57] discovered that heparin can prevent the shedding of the EC GCX in human umbilical vein ECs exposed to plasma samples from COVID-19 patients, and heparin can also restore redox balance by inhibiting the activity of heparanase. Hence, co-treatment of S1P and heparin can protect and synthesize nascent GCX based on the aforementioned studies.

Interestingly, we witnessed *in vitro* that the albumin-bound S1P and heparin co-treatment stimulated WGA, HS, and SDC1 overexpression beyond the untreated atheroprotective UF-levels that were expected to be the threshold. Since degradation of EC GCX is detrimental to overall vascular function and health, we speculate that overexpression of EC GCX could also be harmful and lead to patholigcal disruptions. For instance, an excessively thick EC GCX can impede blood flow and hinder requisite nutrient exchange between circulating blood and surrounding tissues [59,60]. However, no WGA, SDC1, and GPC1 overexpression was observed in the *in vivo* murine model, as the LCA:RCA normalized levels of percent area coverage and thickness were close to 1 in co-treatment conditions. The differing results can be attributed to the discrepancy in co-treatment exposure times: HCAECs were exposed to the co-treatment for 12 hours in dynamic conditions, while mice were administered the co-treatment for 30 minutes prior to data extraction. This suggests that future studies are needed to further understand the pharmacokinetics and pharmacodynamics of the co-treatment and to determine a proper dosage for acute and chronic administration of the therapeutic in patients.

Our study also found that the co-treatment had pleiotropic effects in combating vascular dysfunction and restoring proper vascular functionality via regulating vascular tone and vessel remodeling and reducing vascular permeability. Our initial experiment was focused on examining remodeling at the EC morphology level, a classical experiment. It is well-established that chronic application of UF induces alignment of EC monolayers to the direction of flow and elongation of cell shape [61], along with the planar polarity of intracellular organelles such as the microtubule and the Golgi apparatus [62]. Furthermore, Jung et al. [34] determined that shear-induced EC alignment can be significantly suppressed through S1PR1 downregulation by shRNA or FTY720-P treatment (potent functional antagonist). Jung’s team also determined that the re-expression of S1PR1 restored EC alignment to the direction of flow [34]. In the context of our study, no such improvement of EC alignment with the direction of flow (indicated by the OOP metric) or amongst each other (indicated by the MPA metric) was induced in the atheroprone-DF region after the administration of albumin-S1P and heparan co-treatment. This difference could be due to multiple reasons. Primarily, the results of our experiment performed on 12-hour flow cultured human cells may not be easily comparable to prior published work using ECs from other species which are exposed to flow for a long time period *in vitro* or chronically *in vivo.* Thus, other models such as longer-term EC stimulation by DF and UF may be necessary to maximally align and elongate the ECs before implementation of testing the efficacy of applying albumin-S1P and heparin co-treatment to improve EC alignment in atheroprone-DF regions, and this will be the subject of future work.

When we examined remodeling at the tissue level via ultrasound, we found that vessel wall thickness was substantially reduced in the ligated LCA (DF) in mice administered with co-treatment. Previous studies have also shown the impact that S1P and heparin can individually have on vessel wall thickness. For instance, S1P has been shown to reduce blood pressure by activating S1PR1-mediated release of vasodilatory NO [63]. In pulmonary arterial hypertension, S1P-treated mesenchymal stem cells reduced right ventricular systolic blood pressure and caused a significant reduction in the right ventricular weight ratio and pulmonary vascular wall thickness [64]. Furthermore, heparin can influence vascular wall structure by inhibiting smooth muscle cell proliferation and migration [65–67]. Snow et al. [68] determined that two weeks of heparin treatment prevented intimal thickening and decreased the elastin content in the extracellular matrix domain in the upper and lower arterial intima after rats were subjected to LCA balloon injury. In our study, the combination of albumin-bound S1P and heparin decreased vessel wall thickness in just 30 minutes in ligated LCA, suggesting that the co-treatment is more potent than individual treatments of S1P and heparin. While there was detectable vessel wall thickening and while it could be counteracted on by the albumin-S1P and heparin co-treatment, there were no differences observed in systolic and diastolic vessel diameters throughout all cohorts of mice. Untreated mice did not show vessel luminal narrowing in the LCA within the 5 days after ligation. This can be attributed to this study observing only the early stages of endothelial dysfunction. In contrast to the study by Nam et al. [43], which observed vessel wall thickening in apolipoprotein-E deficient mice on a high-fat diet within one week of surgery, our study used wild-type C57BL/6 mice on a chow diet. This approach aimed to slow the progression of endothelial dysfunction, allowing us to better observe the early effects of the albumin-S1P and heparin co-treatment. Other studies have shown the impact that S1P and heparin individually have on vessel diameter. Katunaric et al. [69] determined that S1P resulted in significant vessel dilation in human arterioles (50 – 200 μm in luminal diameter), except in the presence of S1PR1 inhibitors and when expression of the receptor was reduced. Furthermore, Tangphao et al. [70] witnessed an increase in vasodilation with heparin administration in the dorsal hand vein of human subjects. Therefore, our future studies should be implemented with an appropriate animal model to determine if the co-treatment can increase systolic and diastolic vessel diameters. With this being said, the mechanism of heparin-induced relaxation involves an increased availability of NO related to the local release of histamine [70], prompting us to examine the NO synthesis pathway.

Corroborating the *in vivo* vessel wall thickening data and providing clues related to vessel relaxation and constriction, we witnessed a substantial recovery in p-eNOS expression in atheroprone DF regions in both *in vitro* and *in vivo* conditions following the administration of the co-treatment. This can be accredited to the fact that both S1P and heparin have been individually shown to upregulate p-eNOS and endothelial production of NO. Igarashi et al. [71] provides evidence that S1P treatment in bovine aortic ECs activates Akt, a protein kinase implicated in phosphorylation of eNOS and is mediated by G protein-coupled cell surface receptors (EDG), primarily S1P receptor-1 (S1PR1) on ECs. Thus, S1P can induce eNOS phosphorylation and regulate NO production through the PI3K/Akt pathway [72]. However, S1P-dependent eNOS activation is a double-edged sword. Studies have confirmed that S1P-dependent activation of endothelial S1PR1 receptors promotes vasorelaxation responses and antagonizes vasoconstriction by activating eNOS and the production of NO, even in blood vessels where the overall response to S1P (particularly at high doses) leads to vasoconstriction [73,74]. Hence, the usage of 1 uM of albumin-bound S1P was appropriate since no vasoconstriction (no change in systolic or diastolic diameter) was observed throughout the present *in vivo* studies.

Additional studies have suggested that heparin increases eNOS activity in bovine ECs by a mechanism involving inhibitory guanine nucleotide regulatory protein (Gi) [75,76]. To further elucidate the mechanism of action, Li et al. [77] indicated that interactions between heparin and TMEM184A, a heparin receptor that interacts with and transduces stimulation from heparin in vascular cells, elicited activation of eNOS by increasing its serine 1177 phosphorylation in a calcium-dependent manner. Interestingly, Upchurch et al. [76] discovered that a high dosage of heparin, similar to the levels used in acute cardiovascular treatments, can reduce the production of NO in ECs through a mechanism involving a decrease in steady-state *Nos 3 mRNA* and eNOS protein. This is a similar trend that was observed with S1P. Hence, optimizing the co-treatment concentrations of albumin-bound S1P and heparin is imperative to ensuring proper therapeutic efficacy. The novelty in our work comes from combining the two active biomolecules and observing that the co-treatment improves p-eNOS expression in HCAECs compared to treating HCAECs with either albumin-bound S1P or heparin individually (*see Supplemental* Figure 4).

Interestingly, the co-treatment did not affect total eNOS expression, although an upward trend in total eNOS expression was observed after co-treatment *in vitro* in the atheroprone DF region. Previous studies have shown that total eNOS expression is lower in atheroprone DF regions compared to atheroprotective UF regions [11,78], which is consistent with the current study’s findings of reduced total eNOS expression in DF compared to UF. Harding et al. [11] found that total eNOS was substantially more expressed in the abdominal aorta, where blood flow is UF, than in the inner curvature of the distal aortic arch, where blood flow is DF. They determined that total eNOS exhibited a drastic increase in the abdominal aorta compared to the aortic arch [11]. Although total eNOS expression does not increase in the atheroprone region after co-treatment, our studies found that the proportion of active eNOS is higher in the atheroprone DF region post co-treatment (indicated by increased p-eNOS expression). We are the first to discover that this phenomenon occurs with co-treatment of albumin-bound S1P and heparin.

In addition to regulating vessel remodeling via upregulation of the eNOS vascular tone agent, in atheroprone-DF conditions, the co-treatment of albumin-bound S1P and heparin also improved endothelial barrier function by reducing macrophage uptake within vessel walls and inflammation *in vivo* in atheroprone-DF conditions. These findings can be corroborated with studies that have identified the bioactivity of S1P and heparin individually in reversing vessel hyperpermeability [79]. Lee et al. [30] determined that S1P contributes to activating tight-junction-associated protein zona occludens-1 (ZO-1), which in turn plays an integral role in regulating barrier integrity. ZO-1 is then redistributed to the lamellipodia and cell-cell junctions via the S1PR1/Gi/Akt/Rac pathway, which enhances barrier integrity downstream. Furthermore, Li et al. [80] discovered that heparin reduced human pulmonary microvascular EC permeability induced by lipopolysaccharide and determined that angiopoietin (Ang)/Tie2 signaling pathway represents one of the mechanisms behind how heparin exerts its protective barrier function effect. It is also important to note that both S1P and heparin are potent anti-inflammatory agents which is also corroborated in the present study [81,82].

Upregulation of *EXTL3* encoded protein could explain how the co-treatment can synthesize new HS in atheroprone – DF regions. In this study, we were the first, to our knowledge, to show that in DF regions of the vasculature, downregulation of *EXTL3* was present, which can cause a reduction in HS production by ECs. Marques et al. [83] were able to establish that glycoengineered gastric cancer cells with *EXTL3* gene silencing fully abolished HS expression, further corroborating *EXTL3-*mediated promotion of HS polymerization. We believe that the heparin portion of the co-treatment acts upon this pathway, as *EXTL3* is vital for initiating the synthesis of HS chains [83] increasing GCX expression (particularly GAG HS expression) in atheroprone-DF regions of the vasculature *in vivo*. Additionally, the S1PR1-mediated downstream signaling pathway is a pivotal receptor that the co-treatment could target. We determined that S1PR1 expression was flow-dependent, which can be corroborated by other studies [34,84]. We also determined that the co-treatment in atheroprone-DF regions *in vitro* can increase S1PR1 expression, which is necessary for both acute, mentioned above, and chronic signaling events induced by laminar shear stress in ECs.

Although certain biological pathways can be attributed to the potency of the co-treatment’s ability to protect and synthesize new GCX and improve complex endothelial functions, the mechanical properties of the co-treatment can also play a part in its potency. We found via fluid and particle simulations *in vitro* that longer residence times and higher therapeutic concentrations are present in the atheroprone DF region, where endothelial GCX degradation and dysfunction are most prevalent [8,16,85]. Future studies should look at developing a selective targeting carrier of the co-treatment (i.e. nanoparticle) to further improve the potency and efficacy of the co-treatment in such atheroprone-DF region. Furthermore, future studies should also determine mechanism of action as it pertains to the GCX’s role in maintaining endothelial integrity after co-treatment.

In conclusion, the endothelial GCX is a complex, protein-polysaccharide layer that assists in various endothelial functions including regulation of the barrier at the interface between the blood circulation and the tissue of the blood vessel wall and overall vascular tone health. This structure is prone to mechanical degradation in DF regions. We determined that the co-treatment of albumin-bound S1P and heparin is a potent therapeutic that can protect and synthesize new endothelial GCX. The co-treatment was also able to improve vessel hyperpermeability and vascular tone via increase in eNOS expression and vessel remodeling. This current finding could translate to therapy for preventing atherosclerosis and motivate future development of new therapies targeted at the GCX to treat vascular disease in early stages.

## ABBREVIATIONS

AF: Alexa Fluor
Ang: Angiopoietin
BS: Blocking solution
BSA: Bovine serum albumin
CD68: Cluster of differentiation 68
CILS: Chemical Imaging of Living Systems
DAPI: 4’,6-diamidino-2-phenylindole
DF: Disturbed flow
EC: Endothelial cell
EDG: G protein-coupled cell surface receptors
eNOS: Endothelial-type nitric oxide synthase
*EXTL3*: Exostosin-like glycosyltransferase-3
GA: Glutaraldehyde
GAGs: Glycosaminoglycans
GCX: Glycocalyx
Gi: Inhibitory guanine nucleotide regulatory protein
GPC1: Glypican-1
HCAECs: Human coronary artery endothelial cells
HS: Heparan sulfate
ICAM-1: Intercellular adhesion molecule-1
IV: Intravenous injection
KLF2: Krüppel-like Factor 2
KLF4: Krüppel-like Factor 4
LCA: Left carotid artery
MFI: Mean fluorescent intensity
MPA: Median pairwise alignment
NO: Nitric oxide
NU-IACUC: Northeastern University Institutional Animal Care and Use Committee
OOP: Object orientation parameter
PBS: Phosphate buffered solution
p-eNOS: Phosphorylated endothelial nitric oxide synthase
PFA: Paraformaldehyde
PI3K: Phosphoinositide 3-kinase
RCA: Right carotid artery
S1P: Sphingosine-1-phosphate
S1PR1: S1P receptor-1
SDC1: Syndecan-1
SEM: Standard error of the mean
shRNA: Short hairpin RNA
TNF-α: Tumor necrosis factor-alpha
UF: Uniform flow
VCAM-1: Vascular cell adhesion molecule-1
WGA: Wheat germ agglutinin
ZO-1: Zonula occluden-1

## ACKNOWLEDGEMENTS

Data Availability Statement

The data that support the findings of this study are available in the article and its online supplementary files.

## Author Contributions

RM, KP, SK, ME, and EEE made substantial contributions to the conception/design of this work along with data acquisition, analysis, and interpretation. RM, KP, SK, ME, and EEE drafted this manuscript and reviewed the content critically for intellectual merit. EEE was primarily responsible for funding acquisition, provision of resources, and project supervision. SS, EL, and JO provided additional resources and supervision. All authors assisted in editing the paper. All authors were involved in approval of the final manuscript. All authors agree to be accountable for all aspects of the work in ensuring that questions related to the accuracy or integrity of any part of the work are appropriately investigated and resolved.

## Other Contributors

Thank you to the members of the Ebong Mechanobiology Lab, particularly John Mwangi, Nicholas O’Hare, Mohammad Hamrangsekachaee, and Chinedu Okorafor, for their technical assistance with staining protocols and developing mesh grids for the flow chamber. We also thank our collaborators at Boston Children’s Hospital, Drs. Timothy Hla and Steven Swendeman, for their expertise in S1P. They provided initial protocols for developing albumin-bound S1P and for staining S1PR1, which we then adapted as needed for our model. We are grateful to Dr. Debra Auguste’s Bioresponsive Drug Delivery and Tissue Engineering Lab at Northeastern University, especially Rudolf Abdelmessih, for assistance with the DLS equipment. Additionally, we appreciate the Institute for Chemical Imaging of Living Systems (CILS; RRID:SCR_022681) core facility at Northeastern University for providing confocal imaging (LSM 800) and technical support.

