## Supplemental Figures for "Co-Therapy with S1P and Heparan Sulfate Derivatives to Restore Endothelial Glycocalyx and Combat Pro-Atherosclerotic Endothelial Dysfunction"

Brookhaven Instruments

Basic DLS Report

| Sample | | Results | |
|---|---|---|---|
| Type: | DLS | Eff. Diam. (nm): | 608.38 |
| Sample ID: | Ron_2samples_4 - 2 | Polydispersity: | 0.552 |
| Operator ID: | Unknown Operator | Diffusion Coeff. ($cm^2/s$): | 8.067e-09 |
| SOP ID: | DL_LNEs_RA_Duration 300 s | | |
| Start Date/Time: | 4/19/2023 4:25:57 PM | | |

Notes:

Multimodal Size Distribution:

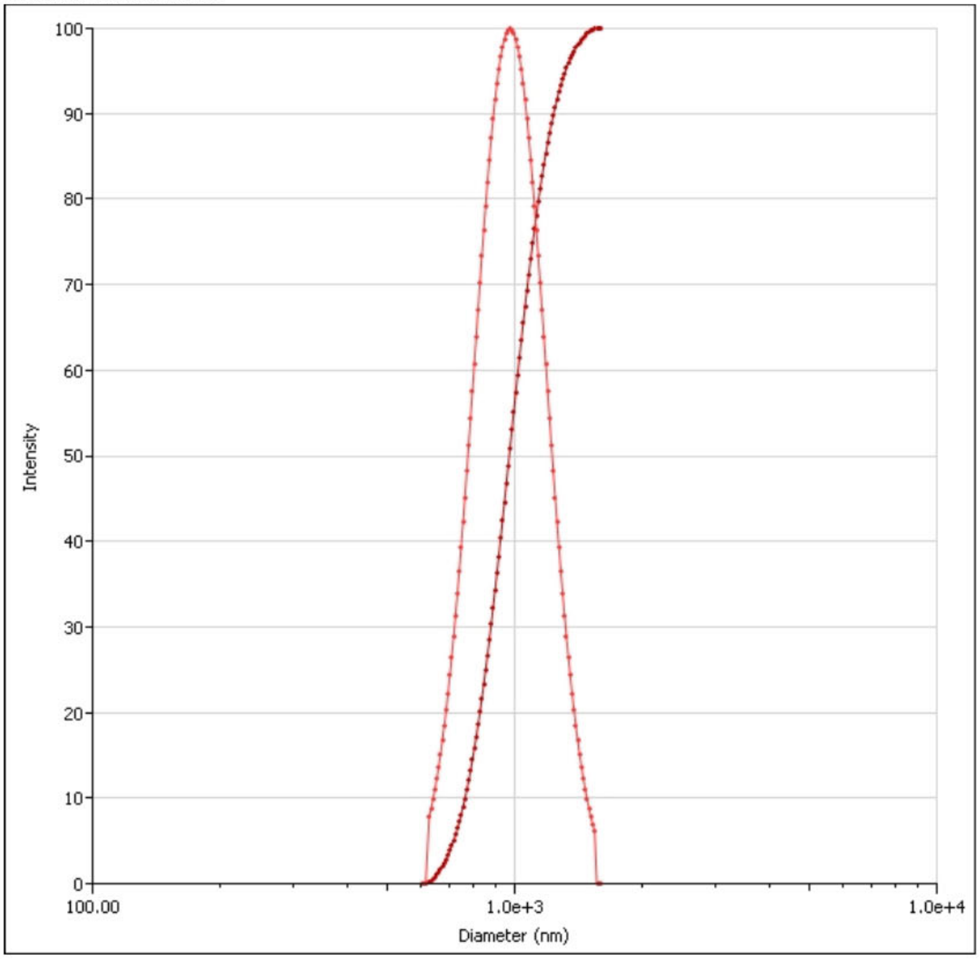


Printed: 4/19/2023 Particle Solutions v. 3.6 (Northeastern University) 2 of 3

***Supplemental Figure 1:***

Dynamic light scattering (DLS) data that was utilized to determine effective diameter (nm) and polydispersity for fluid and particle flow simulations to determine residence time and concentrations in atheroprone – DF region and atheroprotective – UF region of *in vitro* parallel – plate flow chamber.

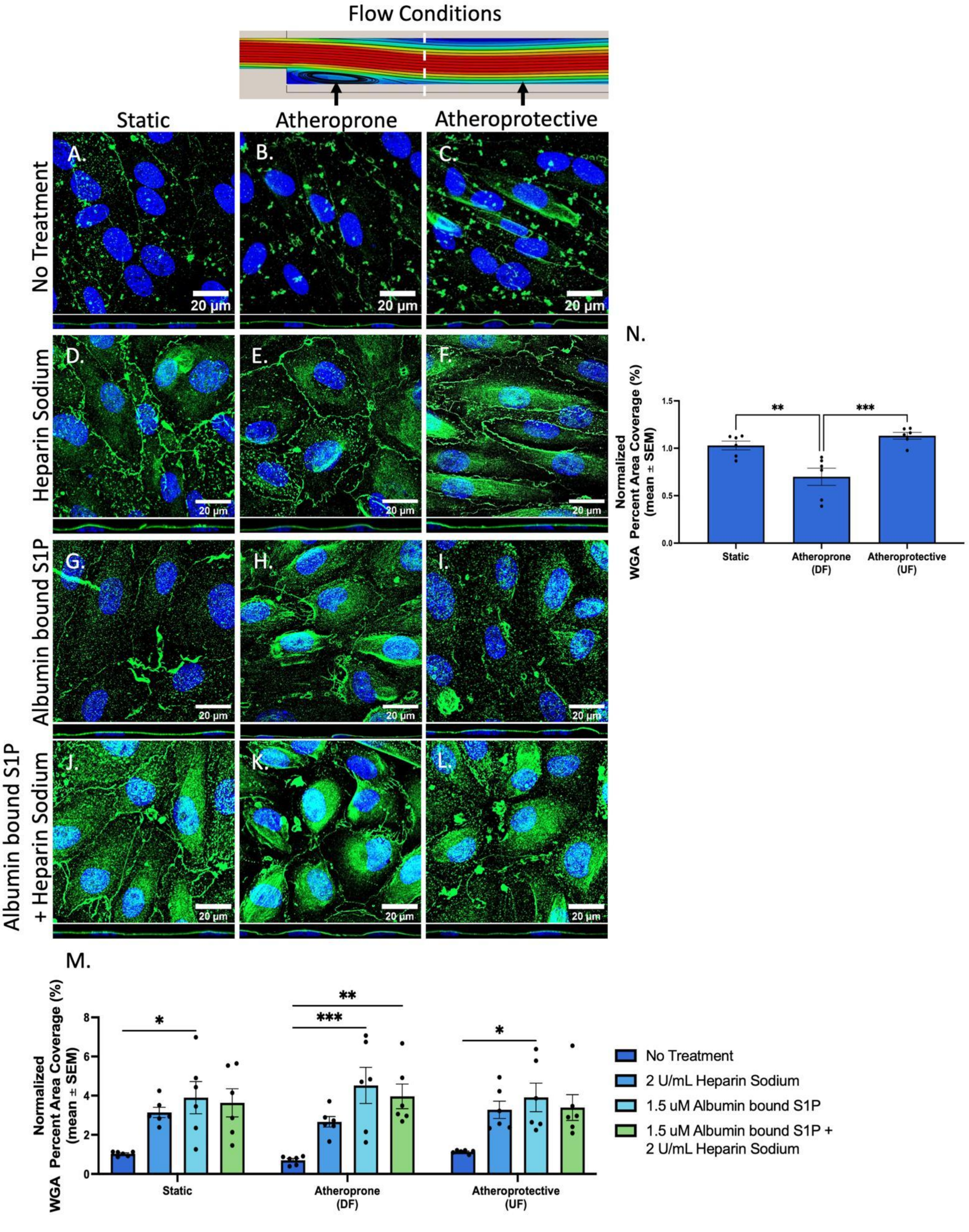

Flow Conditions
Static
Atheroprone
Atheroprotective
No Treatment
Heparin Sodium
Albumin bound S1P
Albumin bound S1P + Heparin Sodium
A.
B.
C.
D.
E.
F.
G.
H.
I.
J.
K.
L.
20 µm
N.
Normalized WGA Percent Area Coverage (%) (mean ± SEM)
Static
Atheroprone (DF)
Atheroprotective (UF)
M.
No Treatment
2 U/mL Heparin Sodium
1.5 uM Albumin bound S1P
1.5 uM Albumin bound S1P + 2 U/mL Heparin Sodium

***Supplemental Figure 2:***

Effect of different treatment groups on wheat germ agglutinin (WGA) expression in HCAECs after 12 hours of flow at 12 dynes/cm$^2$. Illumination of WGA (green), a lectin that stains multiple components of the GCX, was used to provide a comprehensive view and assessment of the percent area coverage of the whole structural GCX. Blue represents cell nuclei labeled with DAPI. The scale bar is 20 μm. These 63x confocal images with oil immersion and z-stacking are representative sections of cells in each group. A) 63x image with oil immersion and z-stacking of HCAECs in the no treatment group under static conditions. B) 63x image with oil immersion and z-stacking of HCAECs in the no treatment group in atheroprone conditions. C) 63x image with oil immersion and z-stacking of HCAECs in the no treatment group under atheroprotective conditions. D) 63x image with oil immersion and z-stacking of HCAECs in the heparin treated group under static conditions. E) 63x image with oil immersion and z-stacking of HCAECs in the heparin treated group in atheroprone conditions. F) 63x image with oil immersion and z-stacking of HCAECs in the heparin treated group in atheroprotective conditions. G) 63x image with oil immersion and z-stacking of HCAECs in the albumin-bound S1P treated group in static conditions. H) 63x image with oil immersion and z-stacking of HCAECs in the albumin-bound S1P treated group in atheroprone flow conditions. I) 63x image with oil immersion and z-stacking of HCAECs in the albumin-bound S1P treated group in atheroprotective flow conditions. J) 63x image with oil immersion and z-stacking of HCAECs in the co-treatment group in static conditions. K) 63x image with oil immersion and z-stacking of HCAECs in the co-treatment group in atheroprone flow conditions. L) 63x image with oil immersion and z-stacking of HCAECs in the co-treatment group in atheroprotective flow conditions. M) Graph shows the mean ± SEM of percent area coverage of WGA staining normalized to the control (0 hr static condition). N) Graph shows the mean ± SEM of percent area coverage of WGA staining normalized to the control (0 hr static condition) of only no treatment group. The mean and error bars representing the SEM are included for each condition with n = 6. Statistical analysis was performed using either one-way or two-way ANOVA. Significance is denoted by asterisks: *p<.05, **p<.01, ***p<.001, and ****p<.0001.

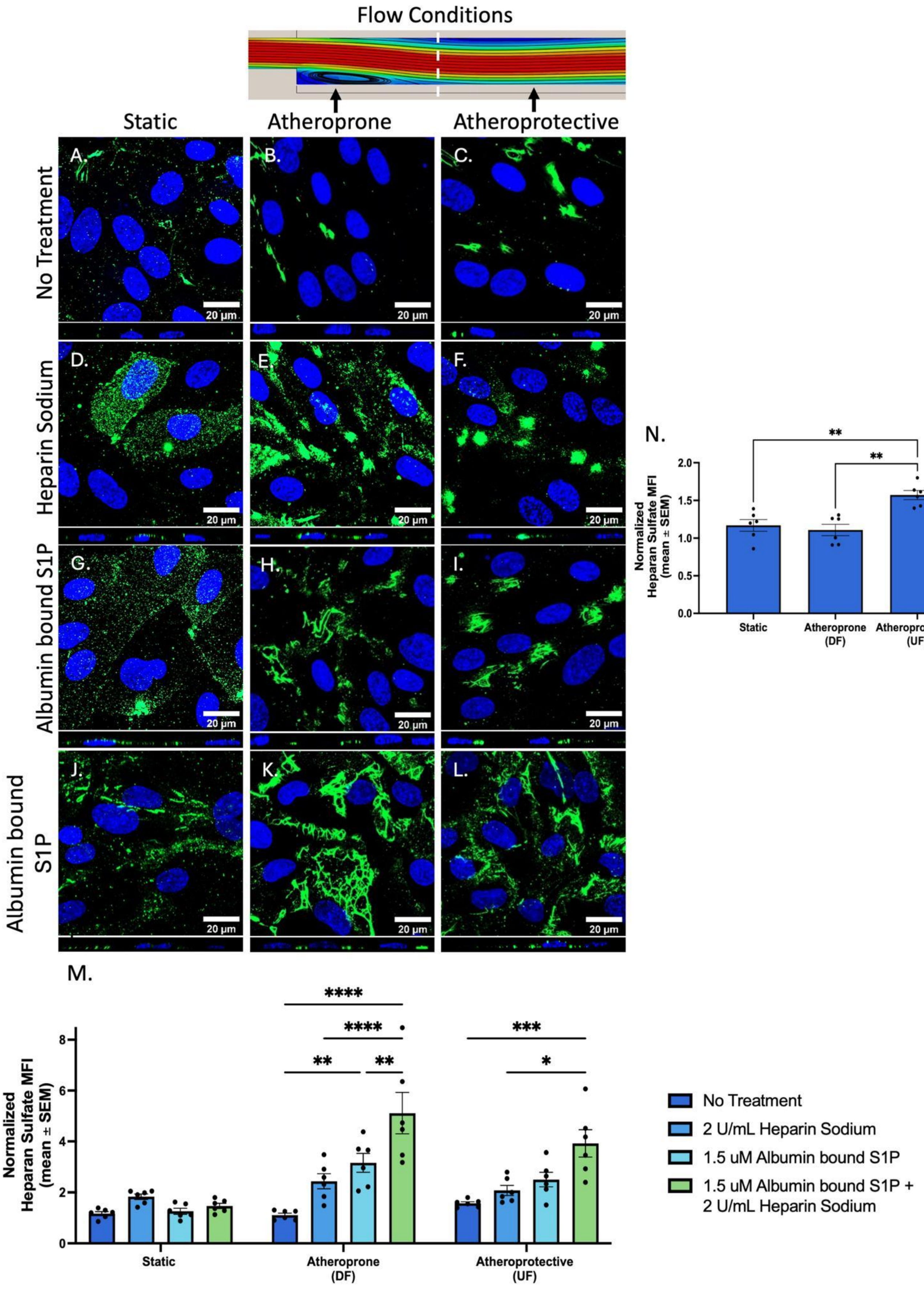

Flow Conditions
Static
Atheroprone
Atheroprotective
No Treatment
Heparin Sodium
Albumin bound S1P
Albumin bound S1P
A.
B.
C.
D.
E.
F.
G.
H.
I.
J.
K.
L.
20 µm
N.
Normalized Heparan Sulfate MFI (mean ± SEM)
2.0
1.5
1.0
0.5
0.0
Static
Atheroprone (DF)
Atheroprotective (UF)
**
**
M.
Normalized Heparan Sulfate MFI (mean ± SEM)
8
6
4
2
0
****
****
**
**
***
*
Static
Atheroprone (DF)
Atheroprotective (UF)
No Treatment
2 U/mL Heparin Sodium
1.5 uM Albumin bound S1P
1.5 uM Albumin bound S1P + 2 U/mL Heparin Sodium

***Supplemental Figure 3:***

Effects of co-treatment on HS expression in HCAECs after 12 hours of flow at 12 dynes/cm$^2$. Illumination of HS (green), a key component of the GCX, was used to assess the mean fluorescent intensity (MFI) of the GCX. The MFI of HS increases in both the atheroprone and atheroprotective conditions when treatment, particularly the co-treatment, is introduced to the flow system. Blue represents cell nuclei labeled with DAPI. The scale bar is 20 µm. These 63x confocal images with oil immersion and z-stacking are representative sections of cells in each group. A) 63x image with oil immersion and z-stacking of HCAECs in the no treatment group under static conditions. B) 63x image with oil immersion and z-stacking of HCAECs in the no treatment group in atheroprone conditions. C) 63x image with oil immersion and z-stacking of HCAECs in the no treatment group under atheroprotective conditions. D) 63x image with oil immersion and z-stacking of HCAECs in the heparin treated group under static conditions. E) 63x image with oil immersion and z-stacking of HCAECs in the heparin treated group in atheroprone conditions. F) 63x image with oil immersion and z-stacking of HCAECs in the heparin treated group in atheroprotective conditions. G) 63x image with oil immersion and z-stacking of HCAECs in the albumin-bound S1P treated group in static conditions. H) 63x image with oil immersion and z-stacking of HCAECs in the albumin-bound S1P treated group in atheroprone flow conditions. I) 63x image with oil immersion and z-stacking of HCAECs in the albumin-bound S1P treated group in atheroprotective flow conditions. J) 63x image with oil immersion and z-stacking of HCAECs in the co-treatment group in static conditions. K) 63x image with oil immersion and z-stacking of HCAECs in the co-treatment group in atheroprone flow conditions. L) 63x image with oil immersion and z-stacking of HCAECs in the co-treatment group in atheroprotective flow conditions. M) Graph shows the mean ± SEM of MFI of HS staining normalized to the control (0 hr static condition). N) Graph shows the mean ± SEM of MFI of HS staining normalized to the control (0 hr static condition) of only no treatment group. The mean and error bars representing the SEM are included for each condition with $n = 6$. Statistical analysis was performed using either one-way or two-way ANOVA. Significance is denoted by asterisks: *$p<.05$, **$p<.01$, ***$p<.001$, and ****$p<.0001$.

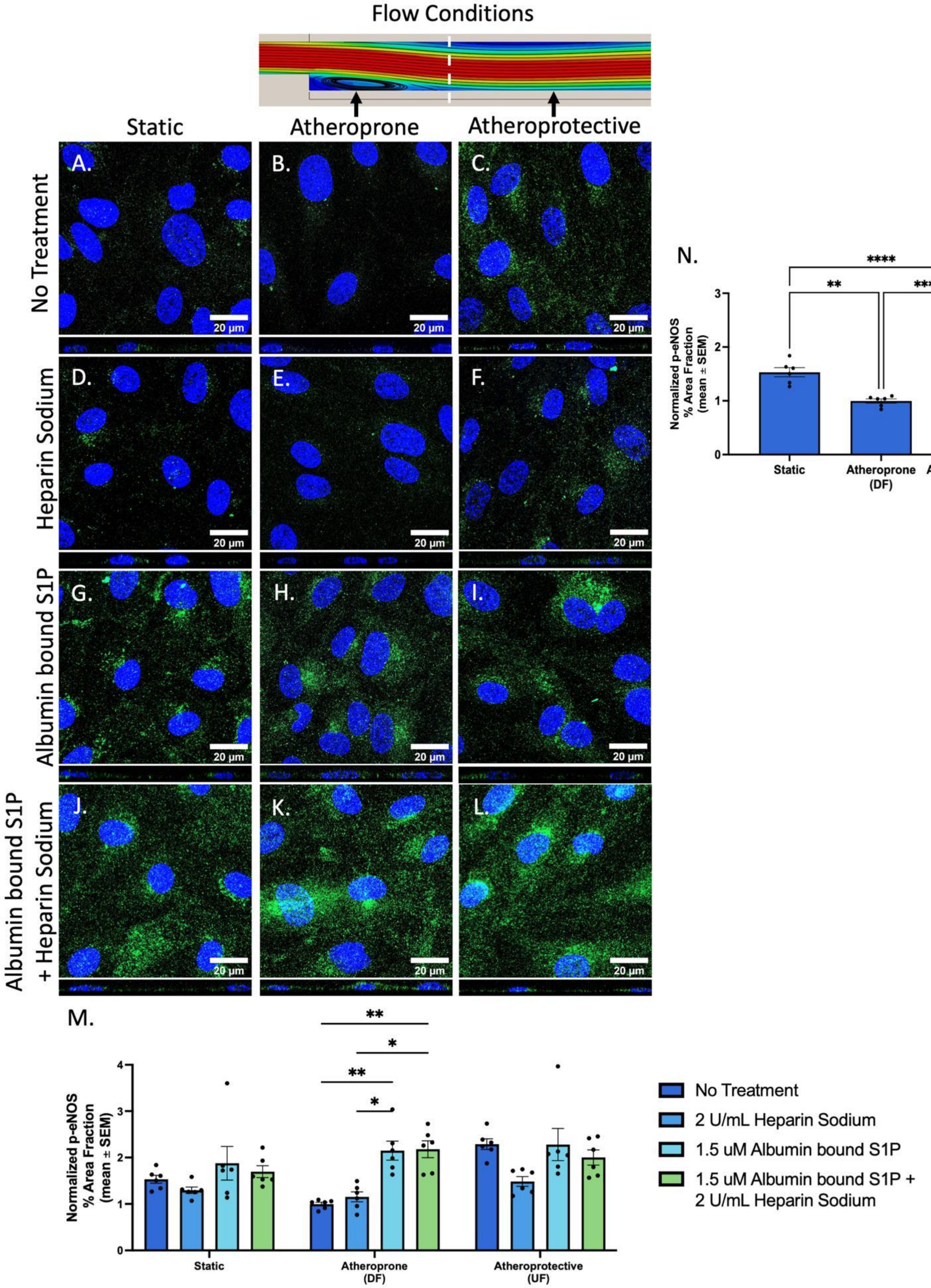

Flow Conditions
Static
Atheroprone
Atheroprotective
No Treatment
Heparin Sodium
Albumin bound S1P
Albumin bound S1P + Heparin Sodium
A.
B.
C.
D.
E.
F.
G.
H.
I.
J.
K.
L.
20 µm
N.
Normalized p-eNOS % Area Fraction (mean ± SEM)
Static
Atheroprone (DF)
Atheroprotective (UF)
**
****
****
M.
Normalized p-eNOS % Area Fraction (mean ± SEM)
Static
Atheroprone (DF)
Atheroprotective (UF)
No Treatment
2 U/mL Heparin Sodium
1.5 uM Albumin bound S1P
1.5 uM Albumin bound S1P + 2 U/mL Heparin Sodium

***Supplemental Figure 4:***

Effect of different treatments on activated eNOS (p-eNOS) expression in HCAECs after 12 hours of flow at 12 dynes/cm$^2$. Illumination of p-eNOS (green), an enzyme that precedes the vasodilator eNO and contributes to vascular tone, was used to assess functional GCX. The percentage area fraction of p-eNOS increases in the atheroprone condition when various treatments, particularly co-treatment, is introduced to the flow system. Blue represents cell nuclei labeled with DAPI. The scale bar is 20 µm. These 63x confocal images with oil immersion and z-stacking are representative images of cells in each group. A) 63x image with oil immersion and z-stacking of HCAECs in the no treatment group under static conditions. B) 63x image with oil immersion and z-stacking of HCAECs in the no treatment group in atheroprone conditions. C) 63x image with oil immersion and z-stacking of HCAECs in the no treatment group under atheroprotective conditions. D) 63x image with oil immersion and z-stacking of HCAECs in the heparin treated group under static conditions. E) 63x image with oil immersion and z-stacking of HCAECs in the heparin treated group in atheroprone conditions. F) 63x image with oil immersion and z-stacking of HCAECs in the heparin treated group in atheroprotective conditions. G) 63x image with oil immersion and z-stacking of HCAECs in the albumin-bound S1P treated group in static conditions. H) 63x image with oil immersion and z-stacking of HCAECs in the albumin-bound S1P treated group in atheroprone flow conditions. I) 63x image with oil immersion and z-stacking of HCAECs in the albumin-bound S1P treated group in atheroprotective flow conditions. J) 63x image with oil immersion and z-stacking of HCAECs in the co-treatment group in static conditions. K) 63x image with oil immersion and z-stacking of HCAECs in the co-treatment group in atheroprone flow conditions. L) 63x image with oil immersion and z-stacking of HCAECs in the co-treatment group in atheroprotective flow conditions. M) Graph shows the mean ± SEM of percent area fraction of p-eNOS normalized to the control (0 hr static condition) for all cohorts. N) Graph shows the mean ± SEM of percent area fraction of p-eNOS normalized to the control (0 hr static condition) of only no treatment group. The mean and error bars representing the SEM are included for each condition with $n = 6$. Statistical analysis was performed using either one-way or two-way ANOVA. Significance is denoted by asterisks: *$p<.05$, **$p<.01$, ***$p<.001$, and ****$p<.0001$.

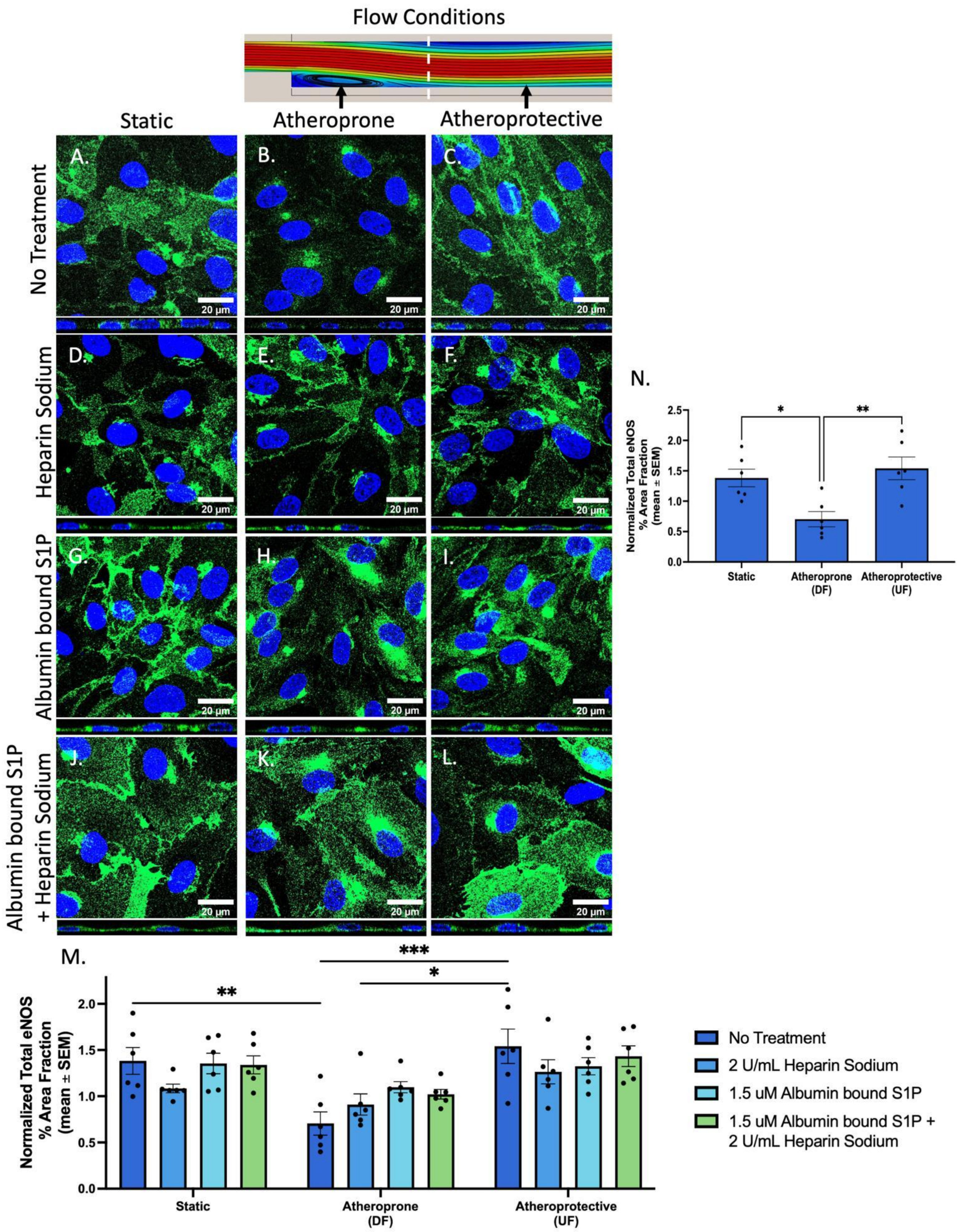

Flow Conditions
Static
Atheroprone
Atheroprotective
No Treatment
Heparin Sodium
Albumin bound S1P
Albumin bound S1P + Heparin Sodium
A.
B.
C.
D.
E.
F.
G.
H.
I.
J.
K.
L.
20 μm
N.
Normalized Total eNOS % Area Fraction (mean ± SEM)
Static
Atheroprone (DF)
Atheroprotective (UF)
M.
No Treatment
2 U/mL Heparin Sodium
1.5 uM Albumin bound S1P
1.5 uM Albumin bound S1P + 2 U/mL Heparin Sodium

***Supplemental Figure 5:***

Effect of various treatments on total eNOS (active and inactive) expression in HCAECs after 12 hours of flow at 12 dynes/cm$^2$. Illumination of total eNOS (green), an enzyme that precedes the vasodilator eNO and contributes to vascular tone, was used to assess functional GCX. No changes in percent area fraction was present in total eNOS expression with and without various treatments. Blue represents cell nuclei labeled with DAPI. The scale bar is 20 µm. These 63x confocal images with oil immersion and z-stacking are representative images of cells in each group. A) 63x image with oil immersion and z-stacking of HCAECs in the no treatment group under static conditions. B) 63x image with oil immersion and z-stacking of HCAECs in the no treatment group in atheroprone conditions. C) 63x image with oil immersion and z-stacking of HCAECs in the no treatment group under atheroprotective conditions. D) 63x image with oil immersion and z-stacking of HCAECs in the heparin treated group under static conditions. E) 63x image with oil immersion and z-stacking of HCAECs in the heparin treated group in atheroprone conditions. F) 63x image with oil immersion and z-stacking of HCAECs in the heparin treated group in atheroprotective conditions. G) 63x image with oil immersion and z-stacking of HCAECs in the albumin-bound S1P treated group in static conditions. H) 63x image with oil immersion and z-stacking of HCAECs in the albumin-bound S1P treated group in atheroprone flow conditions. I) 63x image with oil immersion and z-stacking of HCAECs in the albumin-bound S1P treated group in atheroprotective flow conditions. J) 63x image with oil immersion and z-stacking of HCAECs in the co-treatment group in static conditions. K) 63x image with oil immersion and z-stacking of HCAECs in the co-treatment group in atheroprone flow conditions. L) 63x image with oil immersion and z-stacking of HCAECs in the co-treatment group in atheroprotective flow conditions. M) Graph shows the mean ± SEM of percent area fraction of total eNOS normalized to the control (0 hr static condition) for all cohorts. N) Graph shows the mean ± SEM of percent area fraction of total eNOS normalized to the control (0 hr static condition) of only no treatment group. The mean and error bars representing the SEM are included for each condition with $n = 6$. Statistical analysis was performed using either one-way or two-way ANOVA. Significance is denoted by asterisks: *$p<.05$, **$p<.01$, ***$p<.001$, and ****$p<.0001$.

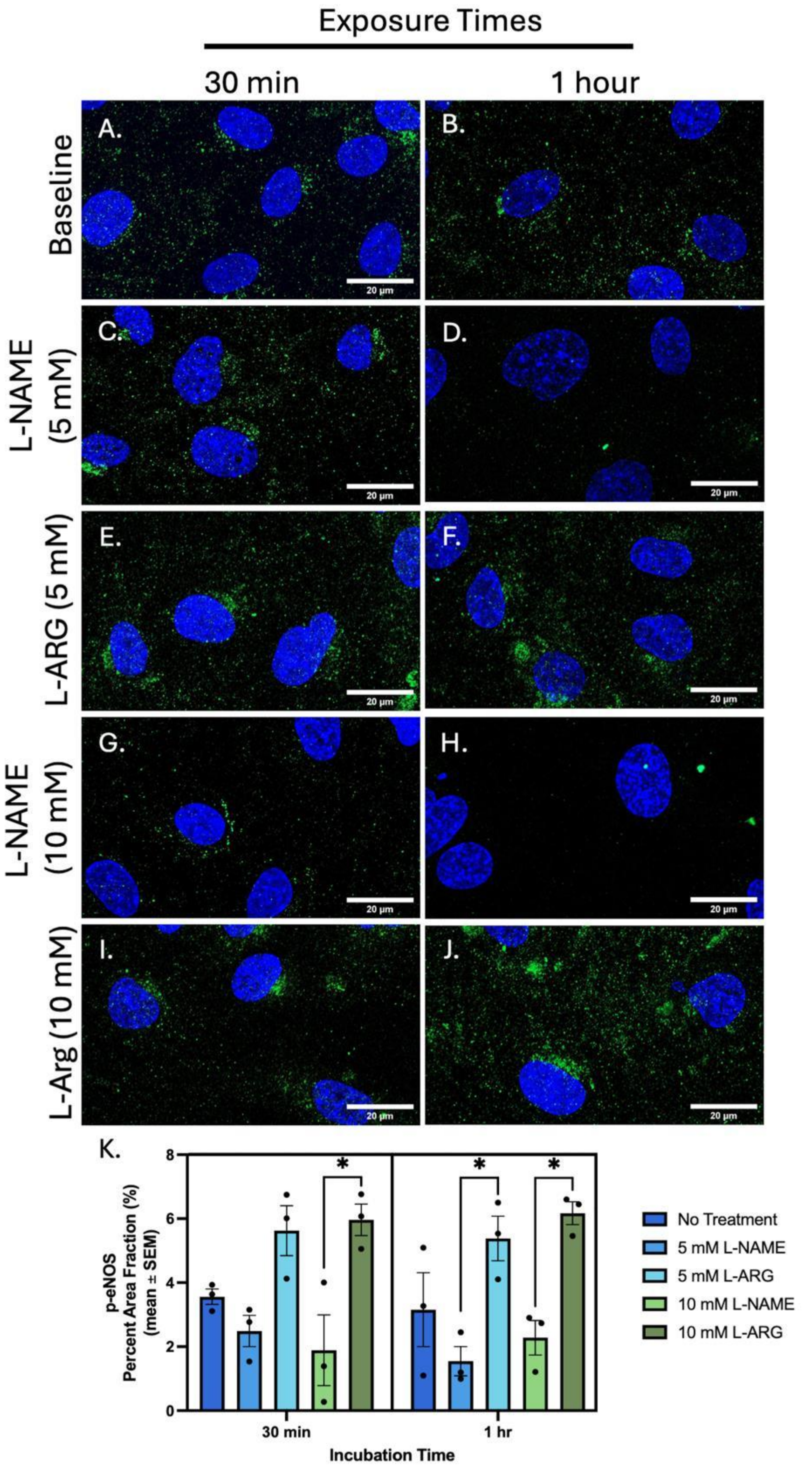
Exposure Times
30 min
1 hour
Baseline
L-NAME (5 mM)
L-ARG (5 mM)
L-NAME (10 mM)
L-Arg (10 mM)
A.
B.
C.
D.
E.
F.
G.
H.
I.
J.
20 μm
K.
p-eNOS
Percent Area Fraction (%)
(mean ± SEM)
0
2
4
6
8
*
30 min
1 hr
Incubation Time
No Treatment
5 mM L-NAME
5 mM L-ARG
10 mM L-NAME
10 mM L-ARG

***Supplemental Figure 6:***

To establish a working immunocytochemistry (ICC) protocol for active eNOS (p-eNOS; green) in HCAECs, a positive control, using L-Arginine ((S)-(+)-Arginine) (L-Arg), which is a potent vasodilator and is the substrate for eNOS to generate nitric oxide (NO). Furthermore, a negative control, using NG-Nitro- L-Arginine Methyl Ester (L-NAME), which is a non-selective NO synthase inhibitor, was also used. As suspected, L-Arg increased p-eNOS expression and L-NAME decreased p-eNOS expression, which suggests a working (ICC) protocol for p-eNOS. Shown above are representative 63x images of HCAECs in all experimental groups. Blue represents cell nuclei labeled with DAPI. The scale bar is 20 µm. (A,B) p-eNOS expression in HCAECs with no L-Arg or L-NAME (baseline). p-eNOS expression in ECs exposed to L-NAME (5 mM) for either C) 30 minutes or D) 1 hr. p-eNOS expression in ECs exposed to L-Arg (5 mM) for either E) 30 minutes or F) 1 hr. p-eNOS expression in ECs exposed to L-NAME (10 mM) for either G) 30 minutes or H) 1 hr. p-eNOS expression in ECs exposed to L-Arg (10 mM) for either I) 30 minutes or J) 1 hr. K) Graph shows the mean ± SEM of percent area fraction of p-eNOS amongst different experimental groups (n = 3). Statistical analysis was performed using two-way ANOVA. Significance is denoted by asterisks: *$p<.05$, **$p<.01$, ***$p<.001$, and ****$p<.0001$.
